# Spatial agent-based modeling and interpretable machine learning identify determinants of combination-therapy response in HER2-heterogeneous breast cancer

**DOI:** 10.64898/2026.03.14.711774

**Authors:** Nizhum Rahman, Trachette L Jackson

## Abstract

HER2 heterogeneity and reversible phenotypic plasticity play a central role in breast cancer progression and therapeutic resistance, yet how their interaction shapes treatment response remains poorly understood. HER2-positive and HER2-negative tumor cell states can dynamically interconvert, enabling compensatory population shifts that undermine monotherapies targeting a single phenotype. Because stochastic lineage effects and local cell interactions are averaged out in mean-field population-level ODE models, we develop a spatially resolved agent-based model (ABM) of heterogeneous tumor growth. We consider paclitaxel, which is modeled to preferentially suppress HER2-positive proliferation, and Notch inhibition, which targets HER2-negative populations and alters phenotypic composition. Starting from single-cell lineages, we assess the ABM against theoretical predictions from a population-level switching model and against single-cell–derived experimental measurements, showing consistency with early lineage dynamics and long-term phenotypic equilibria.

Simulation results show that monotherapies induce compensatory phenotypic shifts and spatial reorganization that permit tumor persistence. In contrast, combination therapy simultaneously targeting HER2-positive and HER2-negative populations disrupts phenotypic replenishment, fragments spatial structure, and can achieve sustained tumor control across a range of simulated tumor regimes and treatment strengths. Importantly, in matched tumors with identical geometry, total burden, and phenotype counts, peripheral enrichment of HER2-negative cells increased post-treatment escape under paclitaxel, whereas this spatial effect was nearly eliminated by combination therapy.

To quantify robustness across heterogeneous tumor parameter regimes, we pair the ABM with an interpretable Random Forest surrogate. Using only pre-treatment and early-trajectory features, the surrogate discriminates sustained control from persistence across held-out simulated parameter regimes and identifies growth-rate asymmetries as dominant drivers of resistance. Together, this integrated mechanistic and data-driven framework clarifies how HER2-mediated plasticity, spatial organization, and competitive growth dynamics shape therapy resistance and provides a scalable approach for analyzing simulated treatment responses across heterogeneous tumor regimes.

## 1 Introduction

HER2 (human epidermal growth factor receptor 2) is a transmembrane tyrosine kinase receptor that plays a pivotal role in the pathogenesis and clinical management of breast cancer. Approximately 15–20% of breast cancers exhibit HER2 gene amplification or protein overexpression, a phenotype associated with aggressive tumor growth, increased recurrence risk, and poor clinical prognosis [1, 2]. HER2 status therefore serves as both a prognostic biomarker and a therapeutic target, guiding the use of HER2-directed agents such as trastuzumab and pertuzumab, as well as newer antibody– drug conjugates [2–5]. Importantly, recent clinical trials have demonstrated therapeutic benefit in tumors with low-level HER2 expression (“HER2-low”), underscoring that HER2 signaling exists along a continuum rather than as a strictly binary classification [5, 6].

While HER2 amplification has traditionally defined a distinct molecular subtype, growing evidence indicates that HER2 expression is dynamic and heterogeneous, particularly in tumors lacking gene amplification. Circulating tumor cells (CTCs) from ER^+^/HER2^−^ patients frequently exhibit HER2^+^ subpopulations during disease progression [7]. These HER2^+^ and HER2^−^ states can interconvert in the absence of new mutations, reflecting reversible phenotypic plasticity that enables adaptation and drug tolerance under therapeutic pressure [7–12].

Clinical observations further suggest that HER2 expression in the absence of gene amplification may be associated with adverse outcomes in otherwise low-risk, node-negative disease that is often excluded from adjuvant anti-HER2 therapy [13, 14]. Together with the emerging “HER2-low” paradigm and the demonstrated activity of trastuzumab deruxtecan in this population, these observations motivate renewed investigation of treatment strategies that explicitly account for HER2-mediated heterogeneity and reversible state transitions [5, 6].

Mathematical models have been proposed to formalize the consequences of phenotypic plasticity in HER2-heterogeneous tumors. In particular, Li and Thirumalai developed a deterministic ordinary differential equation (ODE) model in which HER2^+^ and HER2^−^ populations interconvert and relax to a stable heterogeneous equilibrium, providing a mechanistic explanation for the limited durability of monotherapies targeting a single phenotype [10]. Related population-level models have since extended this framework to incorporate additional ecological interactions and treatment effects [15]. However, well-mixed deterministic models assume homogeneous mixing and large-population averaging. As a result, they neglect spatial constraints, microenvironmental heterogeneity, and stochastic lineage effects that can critically influence clonal expansion and therapeutic escape. Spatially structured mathematical models have been used to investigate how vascular heterogeneity and drug transport influence tumor growth and treatment response in solid tumors [16–18]. In heterogeneous tumors, rare subpopulations and local ecological structure may determine long-term outcome even when mean-field dynamics predict stable coexistence. Spatially resolved agent-based models (ABMs) provide a framework in which individual-cell behaviors, stochastic switching events, and local interactions can be represented explicitly. In this setting, heterogeneous tumor dynamics emerge from biologically interpretable rules rather than population averages [19–21].

Despite these advances, it remains unclear how spatial structure, stochastic lineage effects, and reversible HER2 switching jointly influence treatment response under mono-and combination therapy in heterogeneous tumors. A key unresolved question is whether the spatial localization of a relatively drug-insensitive phenotype can alter treatment response independently of its overall abundance.

Here, we develop a three-dimensional ABM in which individual tumor cells proliferate, die, migrate, and reversibly switch HER2 phenotype. The model incorporates therapeutic interventions with complementary mechanisms of action: paclitaxel, which is modeled to preferentially suppress HER2^+^ proliferation, and Notch inhibition, which targets HER2^−^ populations and modulates phenotypic composition [7, 22, 23]. This framework allows us to study how spatial structure, stochasticity, and phenotypic plasticity jointly shape treatment response under mono-and combination therapy.

To quantify determinants of treatment response across a high-dimensional and nonlinearly interacting parameter space, we pair the agent-based model with a Random Forest (RF) classifier, an ensemble learning method based on bootstrap aggregation of decision trees [24]. This data-driven surrogate is used to discriminate sustained tumor control from persistence across simulated parameter regimes using pre-treatment and early-trajectory features and to identify key drivers of response that complement the underlying mechanistic model. By integrating interpretable machine-learning analysis with spatially resolved simulations, we enable efficient exploration of heterogeneous tumor regimes and identification of determinants of treatment response under the combination-therapy protocol.

Together, our results demonstrate that phenotype localization can alter treatment response even when tumor burden and phenotype abundance are held fixed, revealing a spatial effect that cannot be represented by the corresponding well-mixed population model. Across heterogeneous simulated tumor regimes, the RF analysis further identifies growth-related features as key determinants of sustained response to combination therapy. By combining mechanistic modeling with data-driven analysis, this work provides a unified framework for understanding and analyzing simulated therapy response in HER2-heterogeneous breast cancer.

## 2 Model

We developed a three-dimensional, on-lattice agent-based model (ABM) to investigate intratumoral phenotypic heterogeneity and therapy-driven plasticity in breast tumors with mixed HER2 expression. Individual tumor cells are represented explicitly as agents occupying sites on a cubic lattice, which provides a spatially resolved framework for capturing local cell interactions and stochastic tumor growth.

The tumor population consists of two phenotypic states: HER2^+^ cells (type A) and HER2^−^ cells (type B). These phenotypes differ in their baseline proliferative capacity, death rates, and therapeutic sensitivity. No immune cells, stromal components, or vasculature are included in the present model, allowing the analysis to focus exclusively on tumor-intrinsic phenotypic dynamics.

All tumor-cell behaviors—including proliferation, migration, phenotypic switching, and death—are modeled as stochastic Poisson processes with phenotype-specific rates. Therapeutic interventions, including chemotherapy (Paclitaxel) and targeted therapy (Notch inhibition), are incorporated through time-dependent modifications of these rates over prescribed treatment windows. Model parameters were obtained from the literature when available or selected from biologically reasonable ranges. The baseline parameter values used in the agent-based model are summarized in Table 1, while the treatment-dependent parameter modifications are summarized in Table 2. Complete mathematical update rules and additional parameter details are provided in Appendix A.

**Table 1:** Baseline parameters used in the untreated agent-based model. Event probabilities were implemented using the first-order discrete-time Poisson approximation *p* = *r*Δ*t*.

| Parameter | Description | Value | Source or basis |
| --- | --- | --- | --- |
| $K_1$ | Symmetric division rate of HER2<sup>+</sup> cells | 1.0 week<sup>-1</sup> | Adopted from Li and Thirumalai [10] |
| $K_2$ | Symmetric division rate of HER2<sup>-</sup> cells | 0.7 week<sup>-1</sup> | Adopted from Li and Thirumalai [10] |
| $K_{12}$ | Asymmetric division rate producing a HER2<sup>-</sup> daughter from a HER2<sup>+</sup> parent | 0.09 week<sup>-1</sup> | Adopted from Li and Thirumalai [10] |
| $K_{21}$ | Asymmetric division rate producing a HER2<sup>+</sup> daughter from a HER2<sup>-</sup> parent | 0.09 week<sup>-1</sup> | Adopted from Li and Thirumalai [10] |
| $\delta$ | Baseline death rate of both tumor-cell phenotypes | 0.1 week<sup>-1</sup> | Fixed baseline model assumption |
| $\mu$ | Tumor-cell movement rate | 10 week<sup>-1</sup> | Fixed computational model parameter |
| $\Delta t$ | Simulation time step | 0.1 week | Fixed computational choice |

**Table 2:** Treatment-dependent modifications of phenotype-specific event rates. All values are reported in units of week*^−^*^1^. Parameters not listed as modified retained their baseline values during treatment.

| Parameter | Baseline | During treatment |
| --- | --- | --- |
| *Paclitaxel-modified parameters* |  |  |
| HER2<sup>+</sup> symmetric division rate, $K_1$ | 1.0 | 0.1 |
| HER2<sup>+</sup> asymmetric division rate, $K_{12}$ | 0.09 | 0.27 |
| HER2<sup>+</sup> death rate, $\delta_+$ | 0.1 | 4.0 |
| *Notch-inhibitor-modified parameters* |  |  |
| HER2<sup>-</sup> symmetric division rate, $K_2$ | 0.7 | 0.1 |
| HER2<sup>-</sup> asymmetric division rate, $K_{21}$ | 0.09 | 0.09 |
| HER2<sup>-</sup> death rate, $\delta_-$ | 0.1 | 1.0 |

A schematic overview of the tumor-cell update algorithm is shown in Fig. 1.

**Figure 1:**
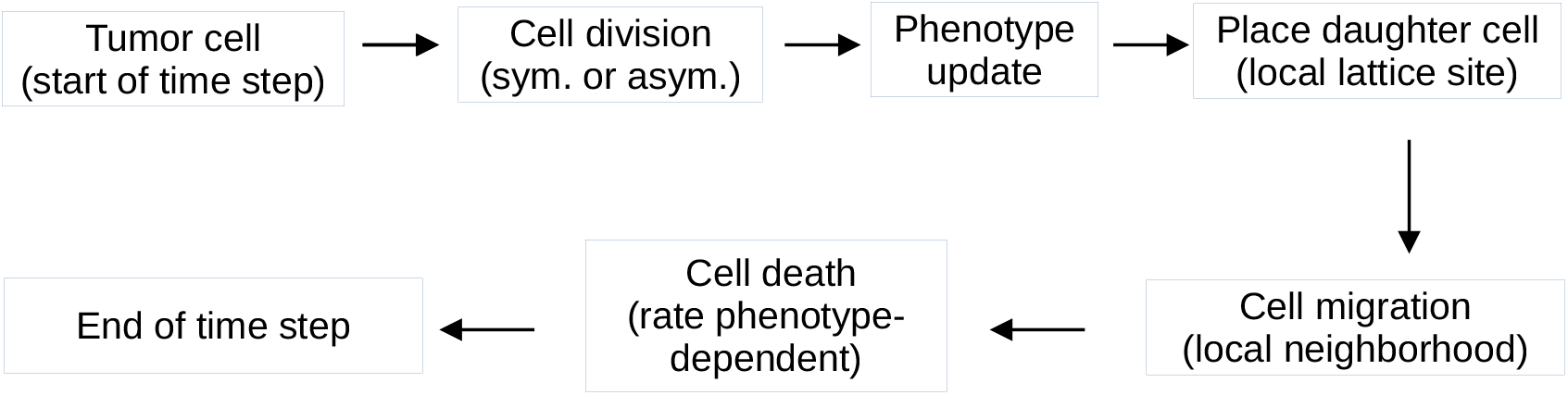
Schematic of the tumor-cell update algorithm in the agent-based model. At each simulation time step, individual tumor cells undergo a sequence of stochastic update events. Cell division is evaluated first and may occur symmetrically or asymmetrically, with phenotype determination coupled to the division event and daughter-cell placement restricted to a neighboring lattice site. Cell migration is then evaluated within the local neighborhood, followed by phenotype-dependent cell death. All event probabilities are governed by phenotype-specific rates. Therapeutic interventions (Paclitaxel and Notch inhibition) are implemented as time-dependent modifiers of these rates and therefore do not alter the update sequence shown.

### 2.1 Tumor cell events

The single-cell lineage and small-colony benchmarking simulations shown in Fig. 2 and Fig. 3 were performed on a 50 *×* 50 *×* 50 cubic lattice, whereas the treatment and spatial-configuration simulations were performed on a 100 *×* 100 *×* 100 cubic lattice. In all cases, non-periodic boundaries were used, such that neighboring sites outside the computational domain were excluded. Both daughter-cell placement and cell migration were restricted to the 26-site three-dimensional Moore neighborhood. If no vacant neighboring site was available following a successful division attempt, no daughter cell was placed; similarly, if no vacant neighboring site was available for migration, the cell remained at its current location. Within each time step, division, migration, and death were applied as separate stochastic attempts, in that order, rather than as mutually exclusive events; thus, a cell could divide and subsequently migrate within the same time step before the death update was applied. Cell order was randomized for migration at each time step, whereas division and death were evaluated according to the lattice/list traversal order used in the implementation.

**Figure 2:**
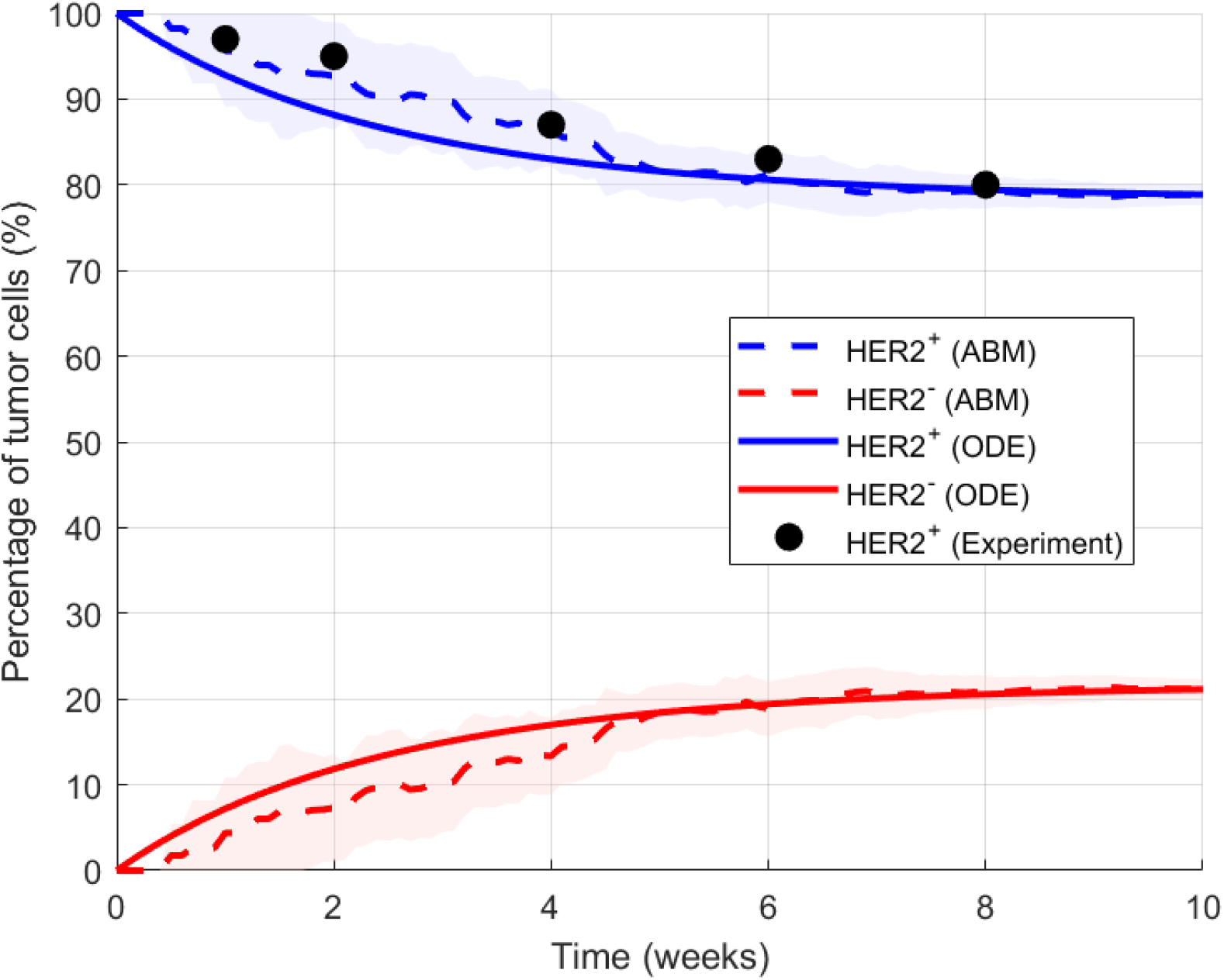
Comparison between the theoretical ODE model, the agent-based model (ABM), and single–cell–derived experimental measurements for HER2^+^ and HER2^−^ fractions. Shaded regions denote 95% confidence intervals from the ABM based on 19 non-extinct stochastic realizations from 20 attempted simulations. Dashed lines show ABM means; solid lines show ODE predictions; filled circles represent experimental HER2^+^ data.

**Figure 3:**
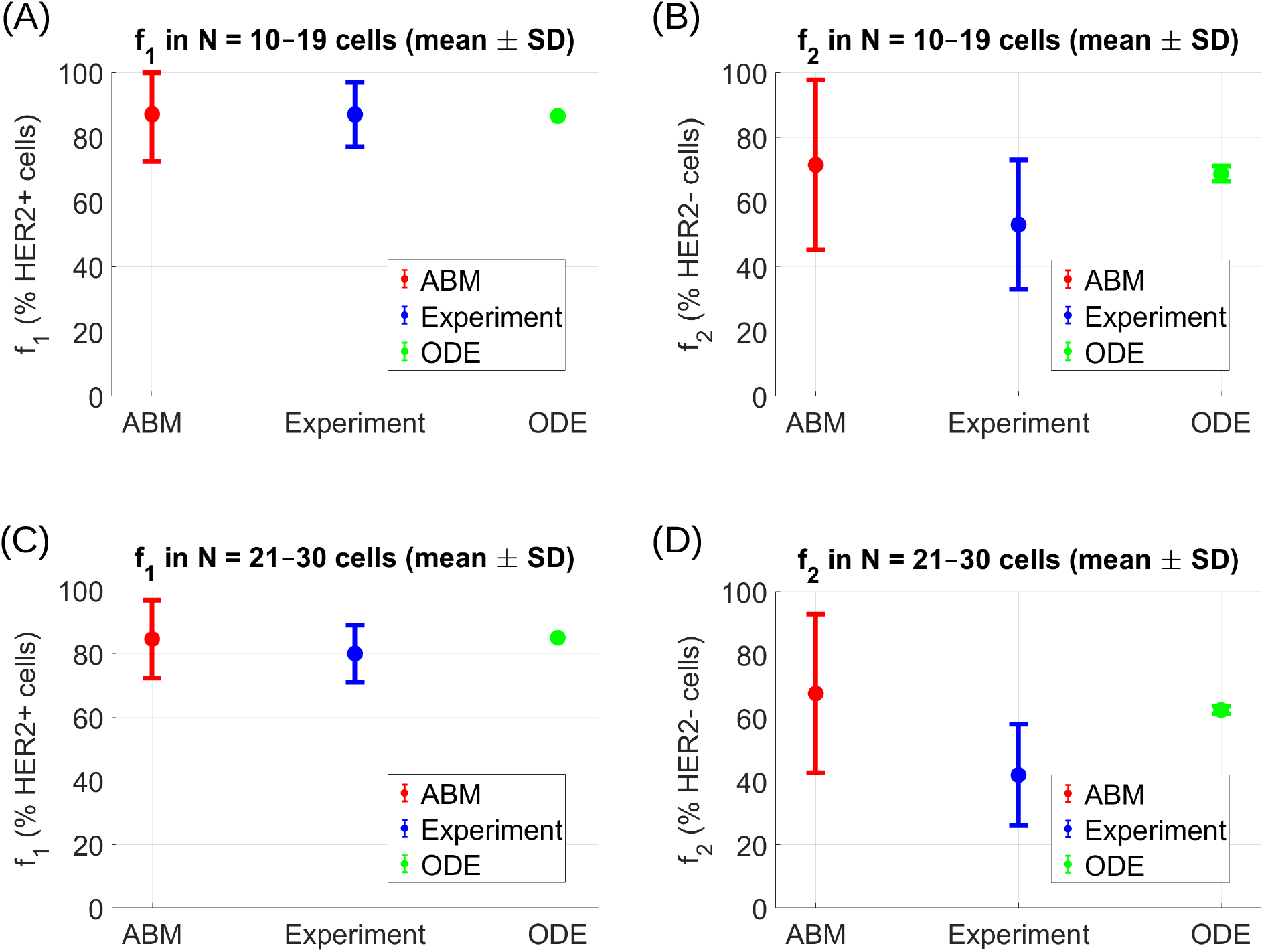
Population-level comparison of HER2 phenotypic fractions predicted by the agent-based model (ABM), experimental measurements, and the deterministic ODE model. Panels (A,C) show HER2^+^ (*f*_1_) fractions for colonies initiated from single HER2^+^ founder cells, whereas panels (B,D) show HER2^−^ (*f*_2_) fractions for colonies initiated from single HER2^−^ founder cells. Panels (A,B) correspond to colonies of size 10–19 cells, while panels (C,D) correspond to colonies of size 21–30 cells. Points denote mean phenotype fractions. For the ABM and experimental data, error bars indicate standard deviations across stochastic realizations or experimental colonies, respectively. For the deterministic ODE, error bars indicate the standard deviation of the predicted phenotype fraction across the integer colony sizes within each specified size interval, rather than stochastic variability.

Cell division may occur symmetrically, producing a daughter cell of the same phenotype as the parent, or asymmetrically, producing a daughter cell of the alternate HER2 phenotype while the parent retains its current phenotype. Phenotypic switching is not implemented as an independent event; rather, it is restricted to division events and is governed by phenotype-specific conversion rates. This choice provides a minimal, lineage-consistent mechanism for reversible state transitions. Future extensions may incorporate continuous or stress-induced switching independent of division to test how non-cell-cycle–coupled plasticity alters treatment response. Cell migration is limited to local lattice neighborhoods, representing short-range motility within dense tumor tissue. Cell death occurs at phenotype-dependent baseline rates.

All event probabilities are computed from their corresponding rates and the simulation time step, ensuring a consistent stochastic interpretation of tumor-cell dynamics.

### 2.2 Paclitaxel treatment

Paclitaxel is modeled as a time-dependent cytotoxic therapy that has stronger effects on the more proliferative HER2^+^ tumor-cell population. This phenotype-conditioned sensitivity is a phenomenological modeling assumption and is not intended to imply HER2-specific molecular targeting by Paclitaxel. During the Paclitaxel treatment window, the division rate of HER2^+^ cells is strongly suppressed, while their death rate is increased to capture chemotherapy-induced cytotoxicity.

In addition to these direct effects, Paclitaxel is assumed to induce phenotypic stress in HER2^+^ cells. This stress is modeled by increasing the rate of asymmetric division from HER2^+^ to HER2^−^, representing therapy-induced transitions toward a more drug-resistant phenotype. All kinetic rates associated with HER2^−^ cells remain unchanged during Paclitaxel exposure. Drug action is applied exclusively over the prescribed treatment interval through a time-dependent control function *u*_tax_(*t*).

### 2.3 Notch inhibitor treatment

Notch inhibition is modeled as a time-dependent targeted therapy that preferentially affects the HER2^−^ tumor-cell population. During the treatment window, the division rate of HER2^−^ cells is reduced and their death rate is increased, representing loss of Notch-dependent proliferative and survival signaling.

Phenotypic plasticity is preserved during Notch inhibition: the rate of asymmetric conversion from HER2^−^ to HER2^+^ remains unchanged. Consequently, Notch inhibition acts through selective pressure on the HER2^−^ compartment rather than by explicitly locking cells into a fixed phenotypic state. All parameters return to baseline values upon treatment withdrawal.

### 2.4 Controlled spatial-configuration experiment

To isolate the effect of phenotype localization on treatment response, we performed a controlled spatial-configuration experiment at the time of therapy initiation. A single untreated tumor was first simulated from week 0 to week 3 using the baseline agent-based model. The resulting week-3 tumor was then used as a common reference configuration for all subsequent spatial comparisons. All three configurations were derived from the same pretreatment tumor realization and therefore had identical occupied geometry, total burden, and global phenotype counts at treatment initiation.

Three matched spatial configurations were generated by redistributing phenotype labels among the occupied lattice sites while preserving the occupied-site mask and the total number of cells of each phenotype. Let **x***_i_* denote the position of occupied lattice site *i*, and let **x***_c_* denote the centroid of all occupied tumor sites. Occupied sites were ranked according to their Euclidean distance ∥**x***_i_* − **x***_c_*∥ from the tumor centroid. For the centrally enriched configuration, HER2*^−^* labels were assigned to the occupied sites closest to the centroid. For the peripherally enriched configuration, HER2*^−^* labels were assigned to the occupied sites farthest from the centroid. For the randomly distributed configuration, the same number of HER2*^−^* labels was assigned to a randomly selected subset of occupied sites. All remaining occupied sites were assigned the HER2^+^ phenotype. The three configurations therefore had identical tumor geometry, total burden, and phenotype counts but differed in their relative spatial organization.

The intended spatial ordering was quantified using an exposed-cell enrichment index. A tumor cell was classified as exposed when at least one site in its 26-site three-dimensional Moore neighborhood was unoccupied. Let *N*_exp_ denote the number of exposed tumor cells, *N*^−^_exp_ the number of exposed HER2*^−^* cells, *N* the total number of tumor cells, and *N ^−^* the total number of HER2*^−^* cells. The exposed-cell enrichment index was defined as

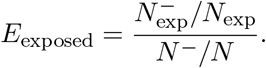

Values of *E*_exposed_ *<* 1, *E*_exposed_ = 1, and *E*_exposed_ *>* 1 indicate depletion, proportional representation, and enrichment of HER2*^−^* cells among exposed tumor sites, respectively.

Each matched spatial configuration was subsequently simulated under three conditions: untreated growth, Paclitaxel monotherapy, and combination therapy consisting of simultaneous Paclitaxel and Notch inhibition. Treatment was applied from week 3 to week 6, and tumors were followed through week 10. For each treatment condition, *n* = 100 stochastic continuations were performed for each spatial configuration. A paired-seed design was used such that continuation replicate *i* used the same random-number seed for the central, random, and peripheral configurations. This pairing enabled direct within-seed comparisons while reducing variation arising from unrelated stochastic realizations. Total tumor burden was normalized to its value at treatment initiation,

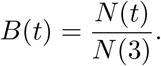

Mean normalized trajectories and pointwise 95% confidence intervals were calculated across the *n* = 100 stochastic replicates. To quantify the effect of HER2*^−^* localization at the final evaluation time, we defined the paired spatial contrast

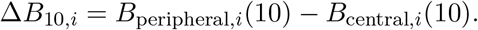

Because corresponding peripheral and central simulations used the same continuation seed, Δ*B*_10*,i*_ provides a within-seed comparison of the two spatial configurations. Positive values indicate greater final tumor burden for peripheral HER2*^−^* enrichment, whereas negative values indicate lower final burden. The mean paired contrast and its 95% confidence interval were summarized across the *n* = 100 matched continuations.

## 3 Results

### 3.1 Single-Cell Lineage Dynamics and Model Benchmarking

Li and Thirumalai [10] developed a deterministic ordinary differential equation (ODE) model to describe population-level dynamics arising from reversible HER2 phenotype switching and to investigate the emergence of cellular heterogeneity. For completeness, we recall the minimal form of their model:

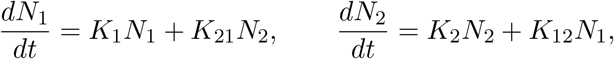

where *N*_1_ and *N*_2_ denote the HER2^+^ and HER2^−^ cell populations, and *K*_1_ and *K*_2_ represent phenotype-specific proliferation rates, while *K*_12_ and *K*_21_ encode reversible switching between phenotypic states. This compact formulation provides a useful theoretical baseline for interpreting lineage-level dynamics driven by phenotypic plasticity.

Our agent-based model (ABM) exhibits behavior that is both qualitatively and quantitatively consistent with the long-term steady state predicted by the ODE model. Starting from a single HER2^+^ founder cell, we simulated the ABM for 10 weeks and tracked the temporal evolution of HER2^+^ and HER2^−^ fractions. Across independent stochastic realizations, the ABM trajectories converge toward a stable phenotypic composition with approximately 78% HER2^+^, in close agreement with the equilibrium fraction predicted by the ODE system.

To further assess the ABM against experimental observations, we compared its predictions with recently reported single–cell–derived lineage measurements. Beginning from a single HER2^+^ founder cell, the experimentally observed HER2^+^ fractions were approximately 97%, 95%, 87%, 83%, and 80% at weeks 1, 2, 4, 6, and 8, respectively. As shown in Fig. 2, these data align more closely with the ABM trajectories than with the deterministic ODE predictions. At these same time points, the mean ABM HER2^+^ percentages were 95.61%, 92.69%, 86.65%, 81.16%, and 79.25%, respectively, whereas the corresponding ODE predictions were 92.80%, 88.16%, 83.00%, 80.59%, and 79.43%. Across the five experimental time points, the mean absolute error was 1.33 percentage points for the ABM, compared with 3.60 percentage points for the deterministic ODE model. In particular, the ABM reproduces the observed pattern of the rapid early decline from a nearly pure HER2^+^ population, whereas the ODE model predicts a smoother and slower relaxation toward steady state. This improved correspondence arises from the ABM’s explicit representation of stochastic single-cell founding effects and early population variability—mechanisms that are averaged out in the mean-field ODE formulation and that influence transient lineage dynamics and early treatment-relevant tumor composition.

### 3.2 Population-Level Experimental Comparisons

Beyond single-cell lineage dynamics, we evaluated whether the ABM reproduces the experimentally observed phenotypic composition of small expanding colonies. Experimental data were obtained from populations initiated from single HER2^+^ or HER2^-^ founder cells and measured after colonies reached sizes within two biologically relevant ranges: 10–19 cells and 21–30 cells. At these small population sizes, stochastic founding effects, asymmetric phenotypic switching, and cell-to-cell variability can produce substantial heterogeneity, providing a useful benchmark for model behavior.

Fig 3 compares the fractions of HER2^+^ and HER2^−^ cells (*f*_1_ and *f*_2_) predicted by the agent-based model (ABM) and the deterministic ODE model against experimental measurements. Panels (A) and (B) correspond to colonies of size 10–19 cells, while panels (C) and (D) show results for colonies of size 21–30 cells. Within each population size, *f*_1_ (HER2^+^) and *f*_2_ (HER2^−^) are shown separately.

For colonies in the 10–19 cell range, the ABM reproduces both the experimental mean and the broad variability observed in HER2 phenotypic composition. For example, the ABM predicts a mean HER2^+^ fraction of 86.50% with a standard deviation of 15.25% (*n* = 100), in close agreement with the experimental mean of 87.00% and standard deviation of 10.00% (*n* = 20) (Fig. 3A). In contrast, the ODE model predicts a nearly identical mean value (86.55%), and the reported ODE standard deviation of 0.63% quantifies variation in the deterministic ODE-predicted phenotype fraction across the integer colony sizes *N* = 10*, . . . ,* 19, rather than stochastic variability.

A similar pattern is observed for the HER2^−^ fraction in the 10–19 cell regime (Fig. 3B). The ABM captures large fluctuations (SD = 26.08%) that closely resemble the experimental variability (SD = 20.00%), whereas the deterministic ODE prediction varies only modestly across the corresponding colony-size interval.

As colony size increases to 21–30 cells, experimental variability decreases due to averaging over a larger number of division events. In this regime, the ODE model predictions move closer to the experimental means, consistent with mean-field behavior, while the ABM continues to capture the observed spread in phenotypic composition (Fig. 3C–D). Together, these results highlight a key modeling principle: the ABM captures stochastic phenotypic variability in small populations, whereas the deterministic ODE system approximates the large-population limit.

### 3.3 Treatment strategy

We next investigated tumor evolution under different therapeutic strategies using a four-panel comparison (Fig. 4). Panel (A) corresponds to untreated growth, while panels (B)–(D) illustrate Paclitaxel monotherapy, Notch inhibitor monotherapy, and combination therapy, respectively. In all treated conditions, therapy was initiated at week 3 and remained continuously active through week 6, after which treatment was withdrawn. Representative two-dimensional spatial slices are shown in Appendix Fig. 14.

**Figure 4:**
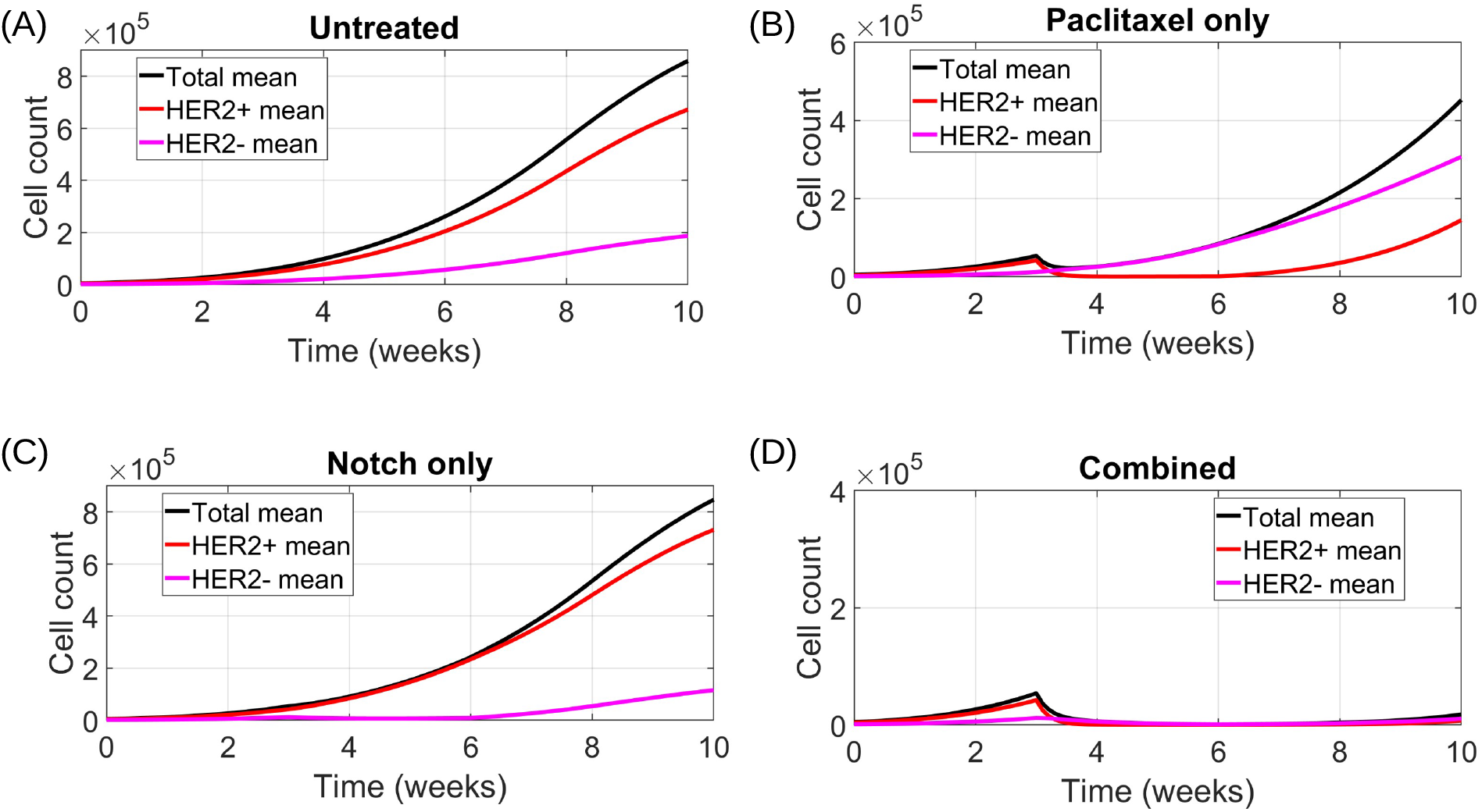
Tumor growth dynamics under different treatment strategies. (A) Untreated. (B) Paclitaxel monotherapy. (C) Notch inhibitor monotherapy. (D) Combination therapy. Black curves denote total tumor burden; red and magenta curves denote HER2^+^ and HER2*^−^* populations, respectively. Curves show means over *n* = 100 independent simulations; 95% confidence intervals are included but narrow.

In the absence of treatment (Fig. 4A), both HER2^+^ and HER2*^−^* populations expand, leading to rapid growth of total tumor burden. Phenotypic coexistence is maintained through ongoing cross-phenotypic conversion.

#### 3.3.1 Paclitaxel monotherapy

Paclitaxel is modeled to preferentially affect the HER2^+^ population by suppressing its proliferative capacity and increasing therapy-induced cell death. As shown in Fig. 4B, Paclitaxel exposure during the treatment interval produces a clear but transient reduction in HER2+ cell numbers, leading to a short-lived decrease in total tumor burden. However, this response is not sustained after treatment withdrawal.

Despite continued Paclitaxel exposure, the HER2^+^ population consistently re-emerges. This rebound is driven by ongoing phenotypic conversion from the HER2*^−^* compartment, which is largely insensitive to Paclitaxel and continues to proliferate during treatment. As HER2*^−^* cells expand, they both replenish the HER2^+^ pool through asymmetric division and progressively dominate the tumor population.

Consequently, Paclitaxel monotherapy induces a shift in phenotypic composition rather than durable tumor control. While HER2+ cells are repeatedly depleted, the untreated HER2– reservoir acts as a source of resistance, enabling tumor regrowth following treatment withdrawal. This compensatory population shift highlights the limitations of single-phenotype targeting in tumors with reversible HER2 plasticity and explains the eventual loss of therapeutic efficacy observed in the population-level dynamics.

#### 3.3.2 Notch inhibitor monotherapy

Notch inhibition selectively targets the HER2*^−^* tumor-cell population by reducing its proliferative capacity and increasing cell death during the treatment window. As shown in Fig. 4C, Notch inhibitor administration produces a marked suppression of HER2– cells during the treatment interval, leading to a transient reduction in their contribution to the total tumor population.

In contrast, the HER2^+^ population remains largely insensitive to Notch inhibition and continues to proliferate throughout treatment. As HER2*^−^* cells are depleted, HER2^+^ cells expand and increasingly dominate the tumor, resulting in sustained overall tumor growth despite effective targeting of the HER2*^−^* compartment. Phenotypic conversion from HER2*^−^* to HER2^+^ further reinforces this trend, allowing the HER2^+^ population to persist even as HER2*^−^* cells are suppressed.

As a result, Notch inhibitor monotherapy alters the phenotypic composition of the tumor without achieving durable control of total tumor burden. These dynamics demonstrate that selectively targeting the HER2*^−^* phenotype alone is insufficient in the presence of reversible HER2 plasticity and an untreated HER2^+^ compartment capable of driving continued tumor expansion.

#### 3.3.3 Combination therapy

Simultaneous administration of Paclitaxel and Notch inhibition produces a qualitatively distinct tumor response compared with either monotherapy. By directly targeting both HER2^+^ and HER2*^−^* populations, combination therapy prevents the compensatory population shifts observed under single-agent treatment. As shown in Fig. 4D, total tumor burden is substantially reduced following treatment initiation and, in many cases, remains suppressed over the simulated time horizon, indicating sustained tumor control.

Unlike Paclitaxel or Notch inhibitor monotherapy, combination therapy limits the ability of one phenotype to expand in response to selective pressure on the other. Continuous depletion of both compartments restricts phenotypic replenishment via asymmetric division, thereby stabilizing tumor dynamics and limiting regrowth following treatment withdrawal.

While this behavior represents the typical response under combination therapy, the ABM also reveals regimes in which dual targeting fails to achieve durable tumor control. Fig 5 shows a representative example for a distinct parameter set under standard-dose combination therapy. Across 100 stochastic realizations, tumor burden initially declines following treatment but subsequently rebounds, resulting in persistent tumor growth. These dynamics demonstrate that combination therapy does not universally guarantee tumor suppression and can fail when intrinsic tumor growth dynamics dominate therapy-induced mortality.

**Figure 5:**
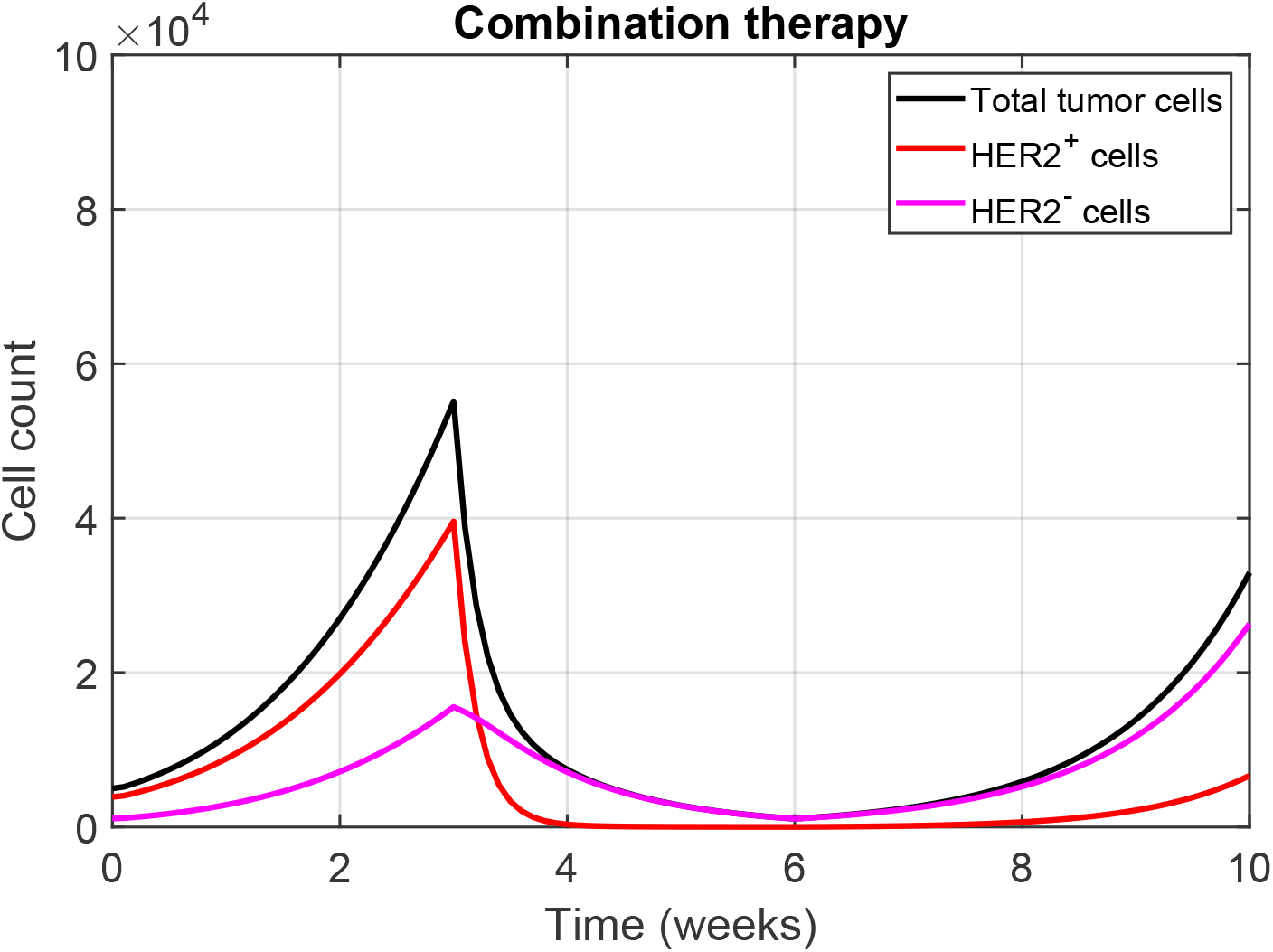
Combination therapy failure under a representative tumor parameter set. Mean tumor trajectories across *n* = 100 stochastic simulations under standard-dose combination therapy. Shaded regions indicate 95% confidence intervals and are narrow relative to the plotting scale. Curves show the total tumor population (black), HER2^+^ cells (red), and HER2*^−^* cells (magenta). Parameter values: *K*_1_ = 0.99, *K*_2_ = 0.89, *K*_12_ = 0.08, and *K*_21_ = 0.05.

To assess robustness with respect to treatment strength, we additionally examined low-and high-dose variants of each therapy. As shown in Appendix Fig. 13, increasing dose primarily enhances tumor control under combination therapy, whereas monotherapies remain insufficient to prevent long-term tumor persistence.

#### 3.3.4 Spatial reorganization under treatment

To complement the population-level dynamics shown in Fig. 4, we next examine the spatial organization of tumor cells under each treatment strategy. The two-dimensional time–space snapshots in Appendix Fig. 14 visualize how tumors reorganize spatially in response to therapy.

Without treatment, the tumor grows outward from the initial cluster in a largely continuous manner, with HER2^+^ and HER2*^−^* cells remaining intermingled as expansion proceeds. Under Paclitaxel treatment, HER2^+^ cells are progressively depleted, and HER2*^−^* cells gradually dominate the spatial structure, filling regions left vacant by therapy-induced cell death. In contrast, Notch inhibition selectively suppresses HER2*^−^* cells, allowing HER2^+^ cells to spread and form a more uniform spatial population.

The most striking behavior occurs under combination therapy. Rather than shrinking uniformly, the tumor fragments into smaller, scattered clusters, and neither phenotype establishes clear spatial dominance. Even at later times, these fragmented clusters persist, indicating that simultaneously targeting both phenotypes disrupts the tumor’s ability to maintain a coherent spatial structure. Overall, these spatial visualizations provide a qualitative view of treatment-induced ecological reorganization that is not visible in well-mixed ODE trajectories. These observations motivated the controlled quantitative spatial analysis described below, in which phenotype localization was varied while tumor geometry, total burden, and phenotype counts were held fixed.

### 3.4 Initial HER2-negative spatial organization alters treatment response

To determine whether phenotype localization itself influences therapeutic response, we performed a controlled spatial-configuration experiment at the time of treatment initiation. A common untreated tumor was first grown to week 3 and then copied into three matched configurations with identical occupied lattice geometry, total tumor burden, and numbers of HER2^+^ and HER2*^−^* cells. The relative spatial organization of the two phenotypes was then altered by assigning the HER2*^−^* population to centrally enriched, randomly distributed, or peripherally enriched locations (Fig. 6A). The corresponding exposed-cell enrichment values were *E*_exposed_ = 0.59, 1.00, and 1.17, respectively, confirming the intended ordering of HER2*^−^* localization. For each treatment condition, we simulated *n* = 100 paired stochastic continuations, using the same continuation seeds across the three spatial configurations.

**Figure 6:**
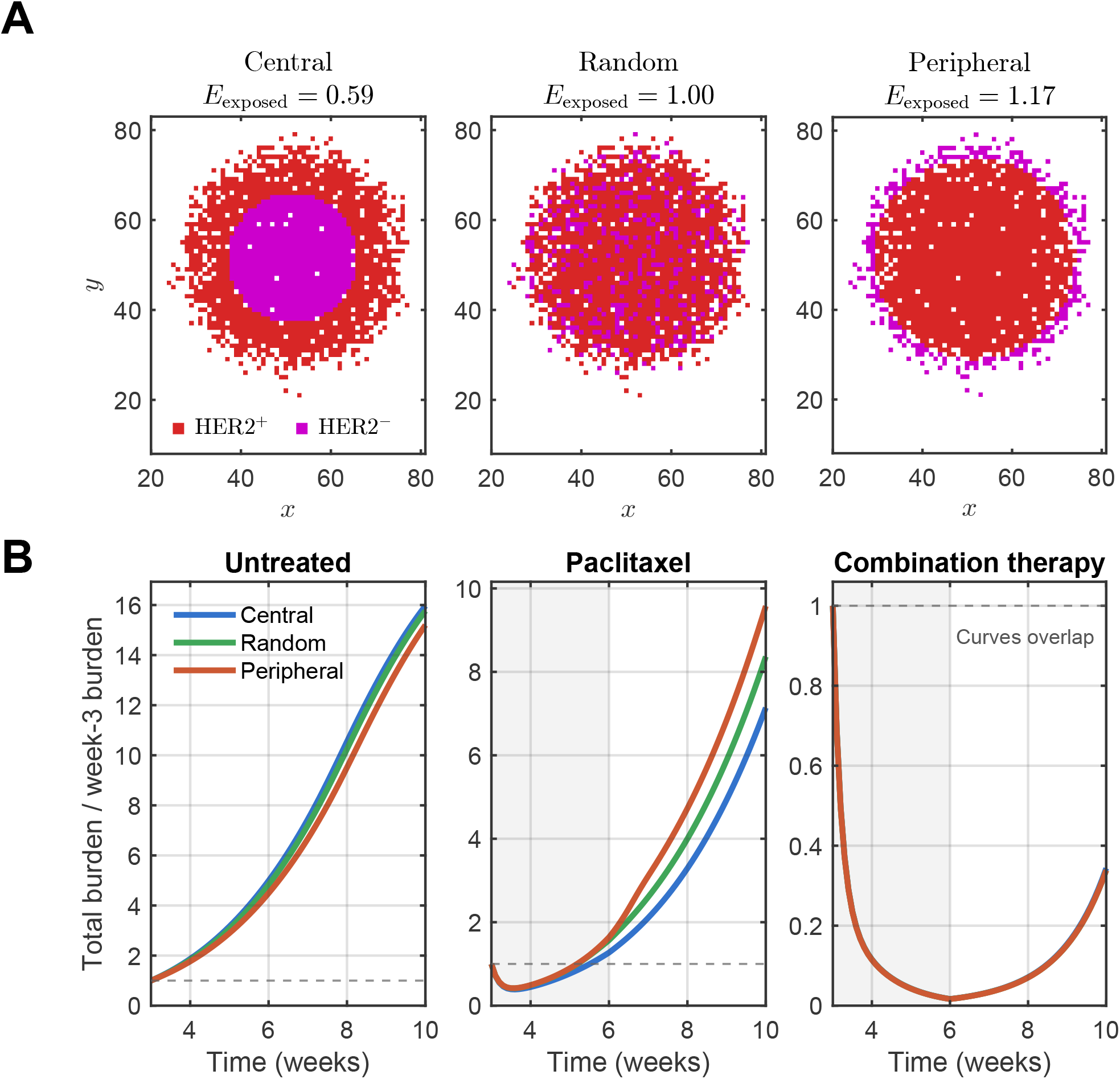
Initial HER2-negative spatial organization alters treatment response. **(A)**Matched tumors at treatment initiation (*t* = 3 weeks) with identical occupied lattice geometry, total tumor burden, and HER2^+^/HER2*^−^* cell counts, but with HER2*^−^* cells arranged in centrally enriched, randomly distributed, or peripherally enriched configurations. Red denotes HER2^+^ cells, magenta denotes HER2*^−^* cells, and white regions denote empty lattice sites. The exposed-cell enrichment index, *E*_exposed_, is defined as the HER2*^−^* fraction among exposed tumor cells divided by the overall HER2*^−^* fraction. Values below, equal to, or above one indicate depletion, proportional representation, or enrichment of HER2*^−^* cells among exposed sites, respectively. **(B)** Mean normalized total tumor burden under untreated growth, Paclitaxel monotherapy, and combination therapy. Tumor burden is normalized by its value at treatment initiation, *B*(*t*) = *N* (*t*)*/N* (3). Solid curves show means across *n* = 100 paired stochastic continuations, and shaded regions indicate pointwise 95% confidence intervals for the means. In several cases, the confidence intervals are narrower than the plotted mean curves. The gray region denotes the treatment interval from weeks 3–6, and the horizontal dashed line denotes the normalized week-3 burden, *B* = 1. The three trajectories under combination therapy overlap almost completely.

In the absence of treatment, all three tumors continued to expand, but their growth trajectories differed modestly according to their initial spatial organization (Fig. 6B). The mean normalized week-10 tumor burdens were 15.969, 15.767, and 15.201 for the centrally enriched, randomly distributed, and peripherally enriched configurations, respectively (Fig. 7A). The paired peripheral-minus-central difference was Δ*B*_10_ = *−*0.768 (95% CI: *−*0.773 to *−*0.763) (Fig. 7B), indicating that peripheral HER2*^−^* enrichment did not provide a general growth advantage in the absence of therapy.

**Figure 7:**
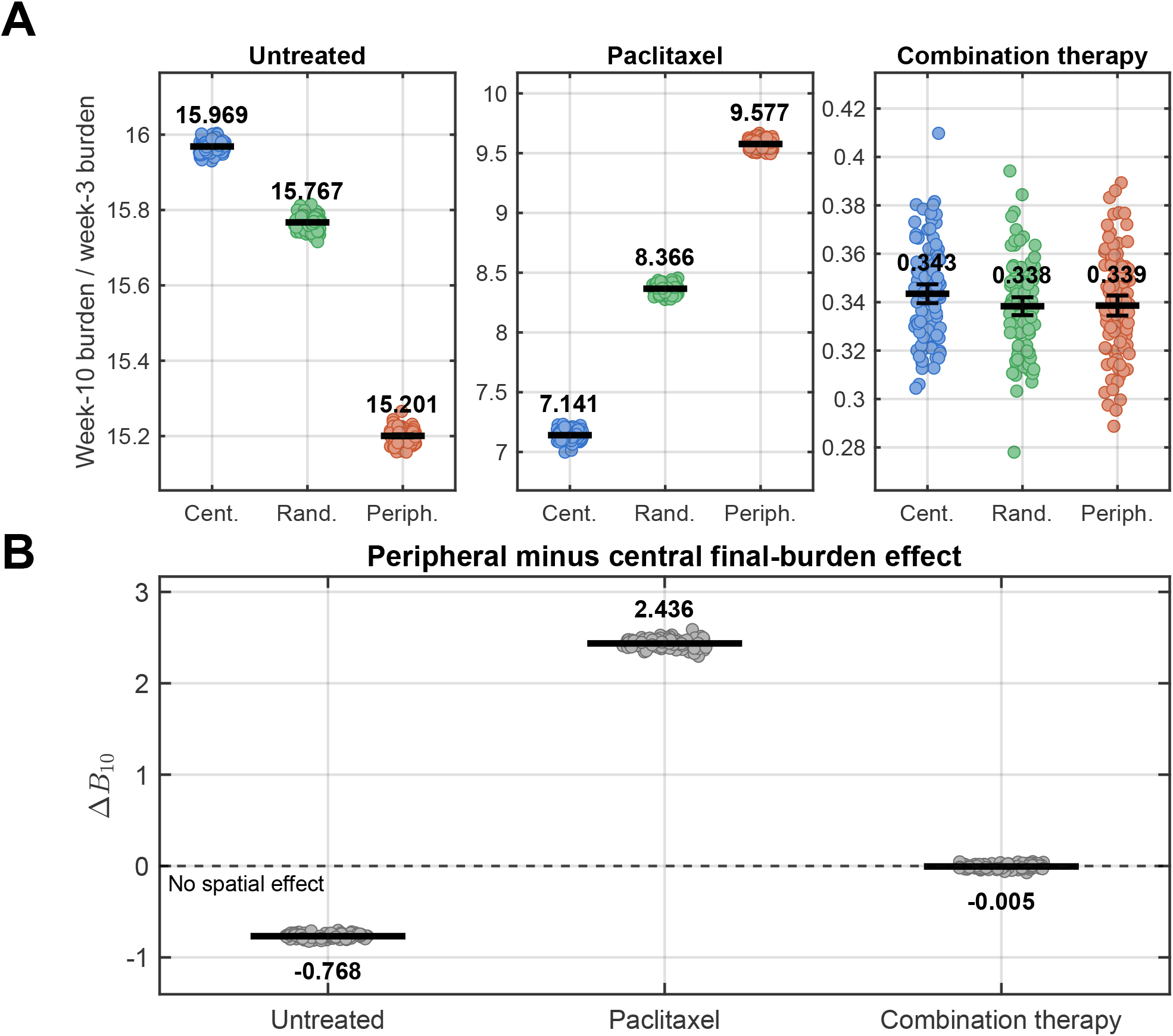
Quantitative effect of initial HER2-negative spatial organization on final tumor burden. **(A)** Individual normalized week-10 tumor burdens for centrally enriched, randomly distributed, and peripherally enriched HER2*^−^* configurations under untreated growth, Paclitaxel monotherapy, and combination therapy. Each point represents one stochastic simulation, thick black horizontal lines denote group means, and error bars indicate 95% confidence intervals for the means. Each spatial configuration contains *n* = 100 simulations. Vertical-axis ranges differ among the treatment panels to display within-treatment variation. **(B)** Paired peripheral-minus-central effect on final normalized tumor burden, defined as Δ*B*_10_ = *B*_peripheral_(10) *− B*_central_(10). Each point represents the paired contrast obtained using the same continuation seed for the peripheral and central configurations. Thick black horizontal lines denote mean paired differences, and error bars indicate 95% confidence intervals for the means. Positive values indicate greater final burden for peripheral HER2*^−^* enrichment, negative values indicate lower final burden, and zero indicates no spatial effect.

A markedly different ordering emerged under Paclitaxel monotherapy. Although all three tumors initially decreased following treatment initiation, the peripherally enriched configuration subsequently exhibited the strongest regrowth (Fig. 6B). Mean normalized week-10 burdens were 7.141 for central enrichment, 8.366 for random distribution, and 9.577 for peripheral enrichment (Fig. 7A). The corresponding paired peripheral-minus-central effect was Δ*B*_10_ = 2.436 (95% CI: 2.427 to 2.445) (Fig. 7B). Thus, relocating the comparatively Paclitaxel-insensitive HER2*^−^* population toward exposed tumor sites substantially increased post-treatment escape. This result is consistent with peripheral HER2*^−^* cells having greater access to locally available space during the selective depletion of HER2^+^ cells.

In contrast, the effect of initial phenotype organization was almost completely eliminated under combination therapy. The mean normalized week-10 burdens were 0.343, 0.338, and 0.339 for the central, random, and peripheral configurations, respectively, and their mean trajectories were nearly indistinguishable (Fig. 6B and Fig. 7A). The paired spatial effect was Δ*B*_10_ = *−*0.005 (95% CI: *−*0.010 to *−*0.0003), indicating a negligible difference in magnitude between the peripherally and centrally enriched tumors (Fig. 7B). Simultaneous targeting of both phenotypic compartments therefore removed the spatial advantage observed for peripheral HER2*^−^* enrichment under Paclitaxel monotherapy.

These treatment-dependent differences represent a specifically spatial prediction of the ABM. At week 3, all three matched tumors contained the same numbers of HER2^+^ and HER2*^−^* cells and therefore corresponded to identical initial conditions in a well-mixed population-level ODE model. Such a model would consequently predict the same treatment trajectory for all three configurations. The distinct Paclitaxel responses observed here arise from phenotype localization, exposed-site access, and local space availability, which are represented explicitly in the spatial ABM but are absent from the ODE formulation.

## 4 Random Forest Analysis

### 4.1 Motivation and overview

We employ a Random Forest (RF) framework to analyze ensemble simulation data generated by the agent-based model (ABM) under *combination therapy* and to assess the robustness of treatment response across heterogeneous tumor parameter regimes. The purpose of this analysis is not comparative prediction across therapy types, but rather to quantify the sensitivity of combination therapy outcomes to intrinsic tumor heterogeneity and early pre-treatment dynamics.

The RF is used as a data-driven surrogate within the sampled parameter domain to identify whether biologically meaningful variation in proliferation, phenotypic switching, and baseline growth leads to differential outcomes under the combined treatment protocol.

#### Primary RF task

We pose the RF analysis as a supervised learning problem in which each ABM simulation replicate yields a feature vector **x** and a binary outcome label *Y* . The task is classification of long-term treatment response under combination therapy at a fixed evaluation horizon *T*_eval_. We define *Y* = 1 as sustained total-tumor control and *Y* = 0 as tumor persistence. Predictions outside the sampled parameter domain are not interpreted mechanistically.

### 4.2 Generation of training data from ABM simulations

Training data for the RF analysis were generated exclusively from ABM simulations of the combination therapy protocol. Intrinsic tumor parameters varied across the ensemble and included phenotype-specific proliferation rates and asymmetric HER2 phenotype switching rates.

A maximin Latin hypercube design was used to sample 200 distinct biological parameter sets. The HER2^+^ proliferation rate was sampled as *K*_1_ *∈* [0.6, 1.0] week*^−^*^1^, while a growth-rate ratio *ρ ∈* [0.3, 1.0] was sampled independently and used to define *K*_2_ = *ρK*_1_. This construction preserved the biological ordering *K*_2_ *≤ K*_1_ while allowing substantial variation in the relative growth rates of the two phenotypes. The asymmetric switching rates were sampled independently as *K*_12_*, K*_21_ *∈* [0.03, 0.15] week*^−^*^1^.

The primary RF dataset used only the standard-dose combination-therapy protocol, with dose scaling fixed at *s* = 1. Treatment was continuously active from week 3 through week 6, and all simulations were followed through week 10. Because treatment dose did not vary within this primary dataset, dose was neither a predictor nor an unobserved source of variation in the RF outcome labels.

For each sampled parameter set, multiple independent stochastic realizations of the ABM were performed to capture run-to-run variability arising from probabilistic cell-level events. Each realization contributed one feature–label pair to the RF dataset.

Ten stochastic realizations were performed for each of the 200 parameter sets, yielding 2,000 simulation-level observations. Before model fitting or inspection of treatment outcomes, 160 parameter sets were assigned to the development partition and 40 parameter sets were reserved as a final held-out test partition. Thus, the development and final-test partitions contained 1,600 and 400 stochastic realizations, respectively. All ten realizations associated with a parameter set remained in the same partition.

### 4.3 Feature definitions and outcome variables

#### No-leakage feature policy

To prevent trivial encoding of treatment outcomes, all RF predictor variables were restricted to information available prior to therapy initiation. Trajectory-derived features were computed over a pre-treatment window [0*, t*_pre_] with *t*_pre_ = 3 weeks. No post-treatment information was used as input to the RF.

#### Predictor variables

Each simulation replicate yields a feature vector **x** capturing intrinsic tumor properties and early dynamics. Predictors include:

- *Intrinsic tumor parameters*, including phenotype-specific proliferation rates and HER2 switching rates (*K*_1_*, K*_2_*, K*_12_*, K*_21_).
- *Pre-treatment dynamic features*, including the effective pre-treatment growth rate *r_b_*, the HER2^+^ fraction at treatment initiation *f*_pre_, and the average pre-treatment change in HER2^+^ fraction *df /dt*_pre_.

The effective pre-treatment growth rate was calculated as

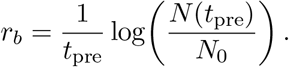

The HER2^+^ fraction at treatment initiation and its average pre-treatment rate of change were calculated as

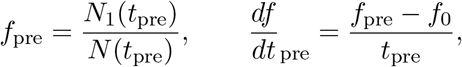

where *f*_0_ = *N*_1_(0)*/N*_0_.

Three RF models were compared: an intrinsic-parameter model using (*K*_1_*, K*_2_*, K*_12_*, K*_21_), a pre-treatment-dynamics model using (*r_b_, f*_pre_*, df /dt*_pre_), and a combined model using all seven predictors.

#### Outcome definition

Let *N* (*t*) denote the total tumor cell count at time *t*. Treatment response was evaluated at *T*_eval_ = 10 weeks.

The primary binary endpoint was defined relative to tumor burden at treatment initiation rather than the single-cell baseline. Sustained total-tumor control (*Y* = 1) was defined as

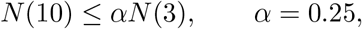

and persistence (*Y* = 0) was defined otherwise. This endpoint corresponds to a reduction of at least 75% relative to the week-3 tumor burden that remains present at week 10, four weeks after treatment withdrawal. Alternative thresholds *α* = 0.10 and *α* = 0.50 were evaluated in a prespecified endpoint sensitivity analysis (Appendix Fig. 15).

### 4.4 ABM outcome structure under combination therapy

Before applying machine-learning analysis, we first summarize the outcome structure generated directly by the agent-based model under combination therapy.

Fig. 8 shows the fractions of stochastic ABM simulations achieving total-tumor, HER2^+^, HER2*^−^*, and dual-phenotype sustained control across low-, standard-, and high-dose combination therapy. For each population, sustained control was defined using the same week-10-to-week-3 threshold *α* = 0.25. This dose-response analysis was generated from a separate exploratory ensemble of 500 ABM simulations in which dose was sampled from *s ∈ {*0.75, 1.00, 1.25*}*. Random allocation of dose produced 163, 165, and 172 simulations at the low, standard, and high dose levels, respectively, explaining the different sample sizes shown in the figure. The dose-response ensemble was used to characterize ABM treatment outcomes, whereas the primary RF analysis was generated separately using only the fixed standard dose *s* = 1.

**Figure 8:**
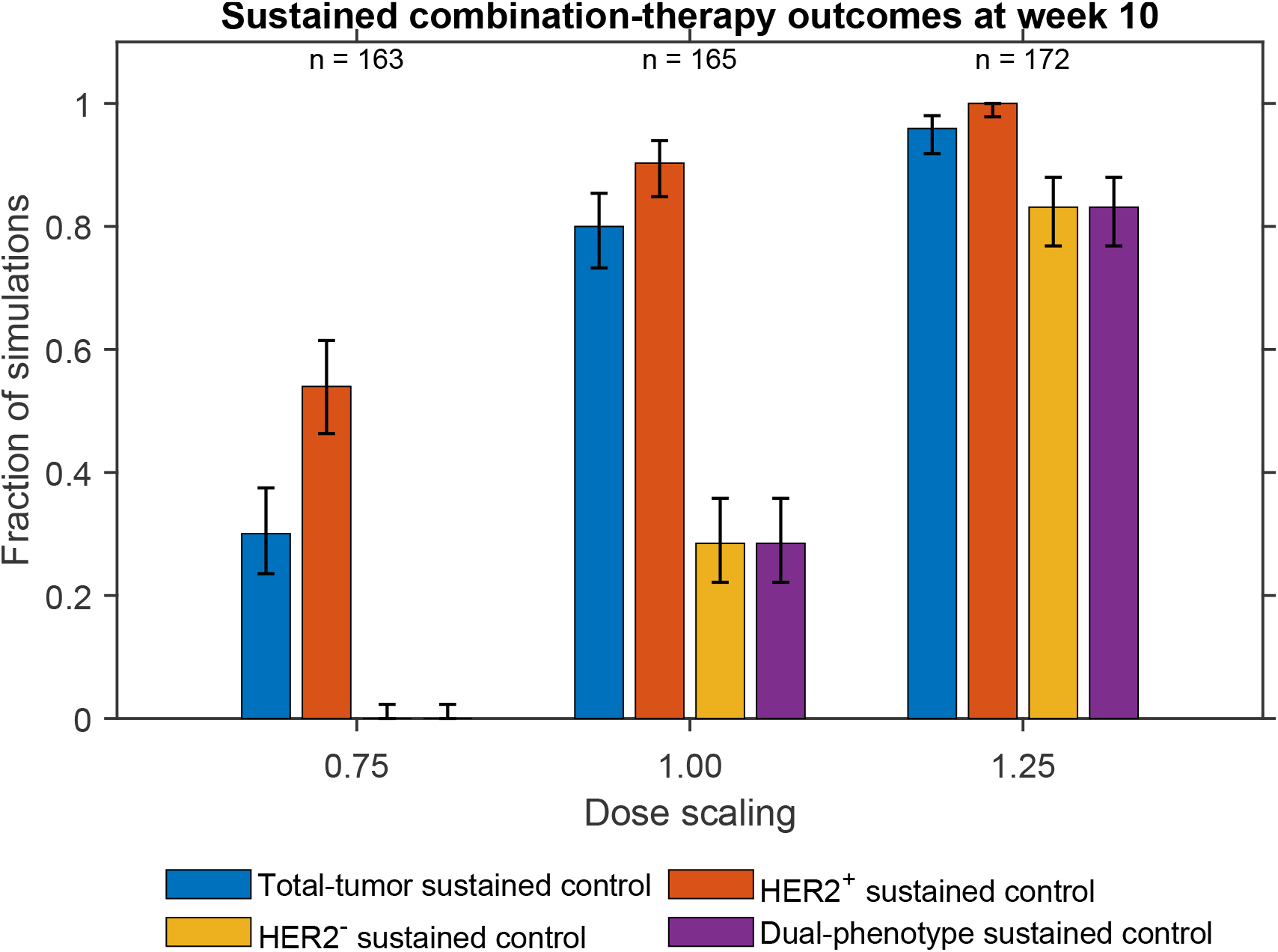
Dose-dependent sustained-control outcomes generated directly by the agent-based model under combination therapy. Bars show the fractions of stochastic ABM simulations achieving total-tumor, HER2^+^, HER2*^−^*, and dual-phenotype sustained control at week 10 across the sampled dose levels. Sustained control was defined as a week-10 population no greater than 25% of the corresponding population at treatment initiation in week 3, representing a sustained reduction of at least 75%. Error bars indicate 95% Wilson binomial confidence intervals, and sample sizes are shown above each dose group. The separate primary Random Forest dataset was generated using only standard-dose combination therapy.

**Figure 9:**
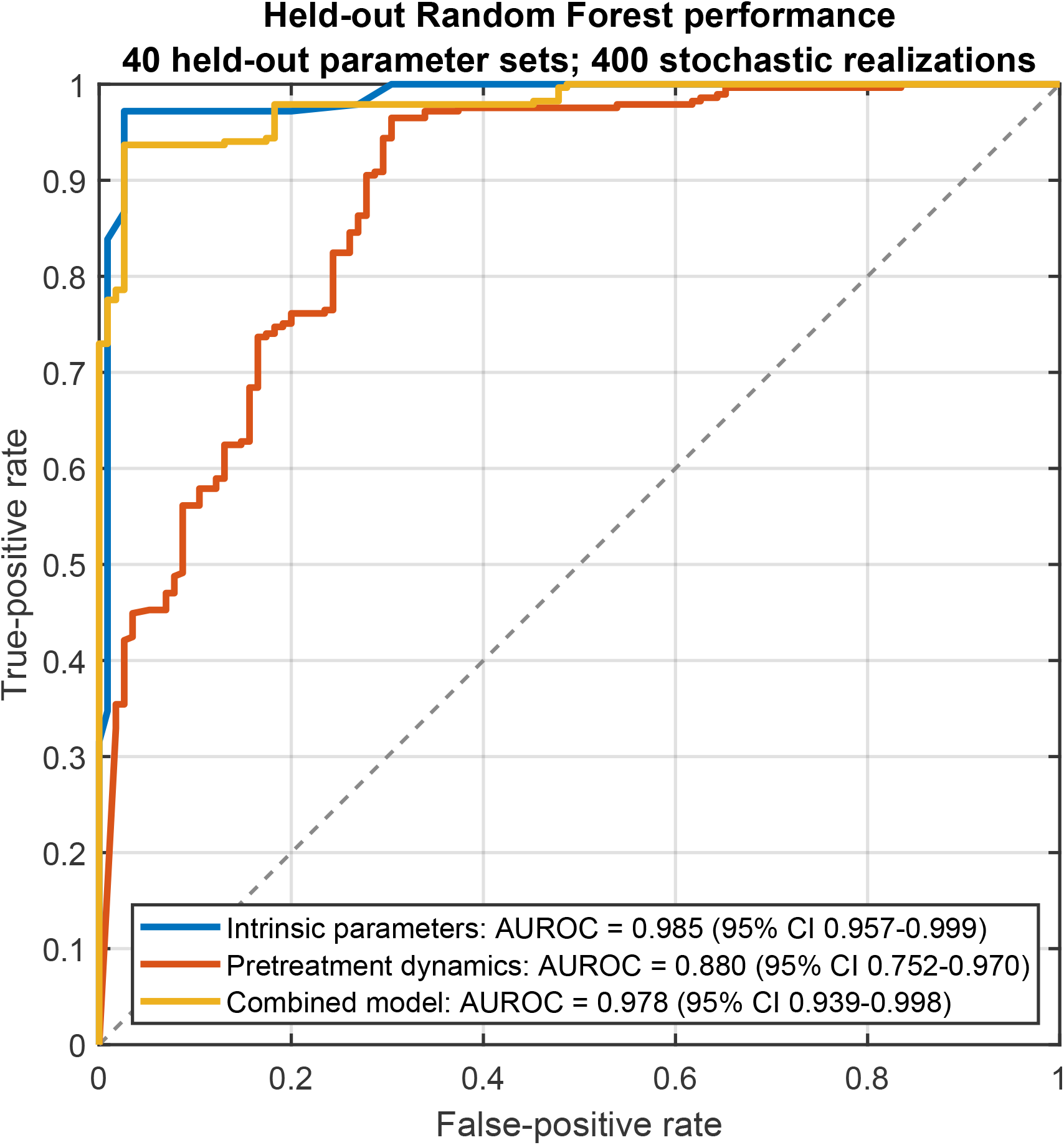
Random Forest generalization to held-out biological parameter regimes under standard-dose combination therapy. Receiver operating characteristic curves are shown for models using intrinsic tumor parameters only, pre-treatment dynamic features only, and the combined predictor set. Models were trained and tuned using 160 development parameter sets and evaluated once on 40 held-out parameter sets comprising 400 stochastic ABM realizations. All realizations associated with a given parameter set were kept within the same partition. The reported 95% confidence intervals for AUROC were obtained by bootstrap resampling complete held-out parameter-set blocks.

### 4.5 Random Forest training and validation

Random Forest classifiers were trained using bootstrap aggregation of decision trees. To ensure evaluation of generalization across biological heterogeneity, data were partitioned using a parameter-wise splitting strategy: all stochastic replicates corresponding to a given parameter set were assigned entirely to either the training or testing partition. This prevents stochastic realizations of the same biological regime from appearing in both partitions and ensures that model performance reflects generalization across distinct parameter combinations rather than memorization of individual simulation runs.

Model development was conducted exclusively using the 160 development parameter sets. Hyperparameters were selected using five-fold grouped cross-validation, with complete parameter sets assigned to individual folds. The minimum terminal-leaf sizes considered were 1, 5, 10, and 20, and the number of candidate predictors considered at each split was tuned separately for each model. Tuning used forests of 300 trees, after which each final RF was refitted using all 160 development parameter sets and 1,000 trees.

Grouped cross-validation selected a minimum terminal-leaf size of 20 and two candidate predictors per split for the intrinsic-parameter model. For both the pre-treatment-dynamics and combined models, the selected minimum terminal-leaf size was 10, with two candidate predictors considered per split. The corresponding grouped development-set AUROCs were 0.985, 0.940, and 0.995 for the intrinsic-parameter, pre-treatment-dynamics, and combined models, respectively.

For threshold-dependent classification, pooled out-of-fold predictions from the development partition were used to select the probability threshold that maximized balanced accuracy. When multiple thresholds produced the same balanced accuracy, the threshold closest to 0.5 was selected. The resulting threshold was then fixed before application to the final-test partition. The final 40 parameter sets were not used for hyperparameter tuning, threshold selection, feature selection, or model fitting. Final-test discrimination was quantified using AUROC. Confidence intervals were calculated by 2,000 bootstrap resamples of complete parameter-set blocks rather than individual stochastic realizations. Balanced accuracy, sensitivity, specificity, precision, F1 score, and average precision were also calculated using the development-selected classification threshold.

On the final held-out test partition, the intrinsic-parameter model achieved an AUROC of 0.985 (95% CI: 0.957–0.999). Using the probability threshold of 0.803 selected exclusively from development-set out-of-fold predictions, this model achieved a balanced accuracy of 0.920 (95% CI: 0.851–0.977), sensitivity of 0.867, specificity of 0.974, precision of 0.988, F1 score of 0.923, and average precision of 0.992 (95% CI: 0.977–0.999). The corresponding confusion matrix contained 247 true positives, 112 true negatives, 3 false positives, and 38 false negatives.

The pre-treatment-dynamics model achieved an AUROC of 0.880 (95% CI: 0.752–0.970). At its development-selected probability threshold of 0.541, the model achieved a balanced accuracy of 0.807 (95% CI: 0.666–0.933), sensitivity of 0.909, specificity of 0.704, precision of 0.884, F1 score of 0.896, and average precision of 0.937 (95% CI: 0.844–0.991). Its confusion matrix contained 259 true positives, 81 true negatives, 34 false positives, and 26 false negatives.

The combined model achieved an AUROC of 0.978 (95% CI: 0.939–0.998). At its development-selected probability threshold of 0.717, it provided the strongest threshold-dependent classification performance, with a balanced accuracy of 0.955 (95% CI: 0.906–0.992), sensitivity of 0.937, specificity of 0.974, precision of 0.989, F1 score of 0.962, and average precision of 0.992 (95% CI: 0.976–0.999). The combined-model confusion matrix contained 267 true positives, 112 true negatives, 3 false positives, and 18 false negatives.

Thus, the intrinsic-parameter and combined models both demonstrated excellent discrimination on previously unseen simulated parameter regimes. Although the intrinsic-parameter model had a marginally higher AUROC, the combined model achieved higher balanced accuracy, sensitivity, and F1 score at its development-selected classification threshold. The pre-treatment-dynamics model remained predictive but showed lower specificity and greater uncertainty than the models containing intrinsic tumor parameters. These results represent internal generalization across simulated parameter regimes within the sampled domain and should not be interpreted as external or clinical patient-level validation.

### 4.6 Sensitivity analysis and interpretation

At the primary threshold *α* = 0.25, 1,430 of the 2,000 simulations (71.5%) achieved sustained total-tumor control. The final-test partition contained 285 controlled and 115 persistent outcomes. Repeating the RF analysis using *α* = 0.10, 0.25, and 0.50 showed that the intrinsic and combined models retained high held-out discrimination across the alternative endpoint definitions. The pre-treatment-dynamics model remained predictive but exhibited greater variation across thresholds. The corresponding endpoint-threshold sensitivity analysis is shown in Appendix Fig. 15.

To identify which biological variables most strongly influence combination therapy outcomes, we quantified feature importance using permutation importance.

Feature importance was calculated using the fitted combined model and the untouched final-test partition. For each predictor, complete ten-realization feature blocks were reassigned among held-out parameter sets, and importance was quantified as the resulting decrease in held-out AUROC. This grouped procedure preserved the within-parameter-set stochastic structure and avoided the replicate-level leakage that could arise from conventional out-of-bag importance. A total of 1,000 grouped permutations was performed for each predictor.

The HER2*^−^* proliferation rate *K*_2_ produced by far the largest decrease in held-out AUROC when permuted, identifying it as the dominant predictor used by the combined RF (Fig. 10). The effective pre-treatment growth rate *r_b_*, the pre-treatment change in HER2^+^ fraction, and *K*_21_ produced substantially smaller reductions, while the remaining variables contributed little additional held-out discrimination. Because the sampling design imposed *K*_2_ = *ρK*_1_ and several predictors are statistically related, individual permutation scores represent the fitted model’s reliance on each variable rather than independent causal effects. The comparatively modest contributions of *K*_12_ and *K*_21_ support interpretation of phenotype switching as a secondary, modulatory influence under the fixed combination-therapy protocol rather than as the principal determinant of escape.

**Figure 10:**
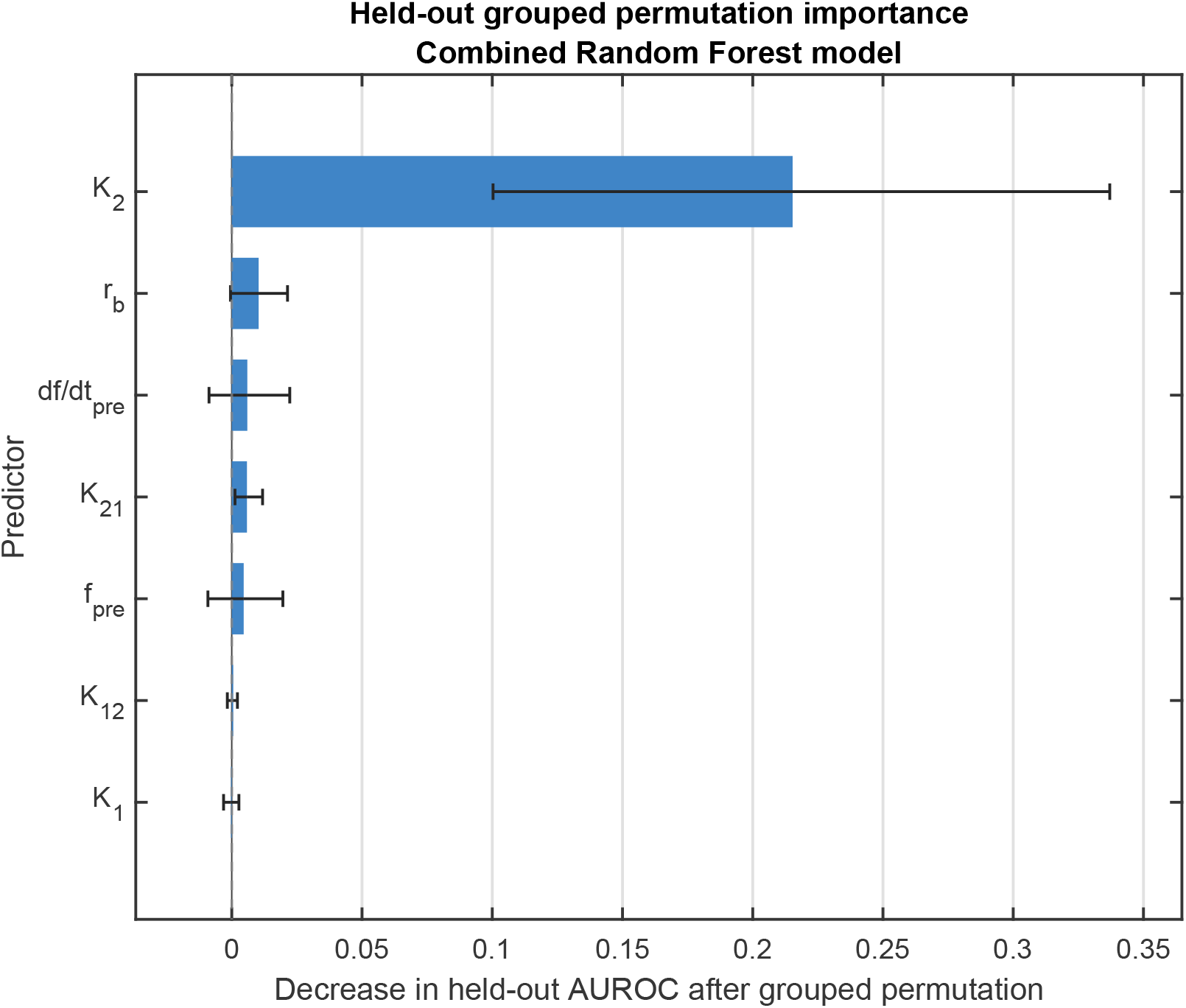
Held-out grouped permutation importance for the combined Random Forest model under standard-dose combination therapy. Importance was measured as the decrease in held-out AUROC after permuting each predictor across complete parameter-set blocks. This procedure preserves the ten stochastic realizations associated with each biological parameter regime and avoids the replicate-level leakage that can occur with conventional out-of-bag importance. Bars show mean decreases in AUROC across 1,000 grouped permutations, and error bars show the corresponding 95% permutation intervals. Larger decreases indicate greater reliance of the fitted model on that predictor.

### 4.7 Link to mechanistic modeling

The RF analysis complements the ABM by formalizing how combination therapy outcomes vary across heterogeneous tumor parameter regimes. Consistent with the ABM simulations, the RF identifies intrinsic tumor growth parameters and early pre-treatment expansion dynamics as dominant drivers of therapeutic success or failure, with phenotype switching rates playing a secondary, modulatory role.

Together, these results demonstrate that the RF learns a compact, interpretable surrogate of ABM-generated outcomes that interpolates across previously unseen parameter combinations within the sampled domain.

### 4.8 Growth-dominated control under combination therapy

While permutation-based feature importance identifies which variables dominate outcome classification, it does not reveal how variations in these parameters shape the treatment response landscape.

To visualize the joint influence of the leading intrinsic and pre-treatment predictors, we evaluated a conditional RF response surface as a function of *K*_2_ and *r_b_*. Predictions were generated using the final combined model. Non-plotted predictors were fixed at their development-set median values, except that *K*_1_ was adjusted with *K*_2_ using the median sampled *K*_2_*/K*_1_ ratio so that the plotted combinations remained consistent with the sampling constraint *K*_2_ *≤ K*_1_.

Fig. 11 shows a pronounced transition across the *K*_2_–*r_b_* response landscape. Lower HER2*^−^* pro-liferation and slower pre-treatment expansion are associated with high predicted probabilities of sustained total-tumor control, whereas increasing either growth-related quantity shifts the model toward treatment escape. This structure indicates that baseline competitive fitness and net pre-treatment expansion largely determine whether standard-dose combination therapy can maintain long-term tumor control. Together with the grouped permutation analysis, the response surface supports a growth-dominated interpretation of treatment resistance, with phenotype-switching parameters contributing comparatively smaller modulatory effects.

**Figure 11:**
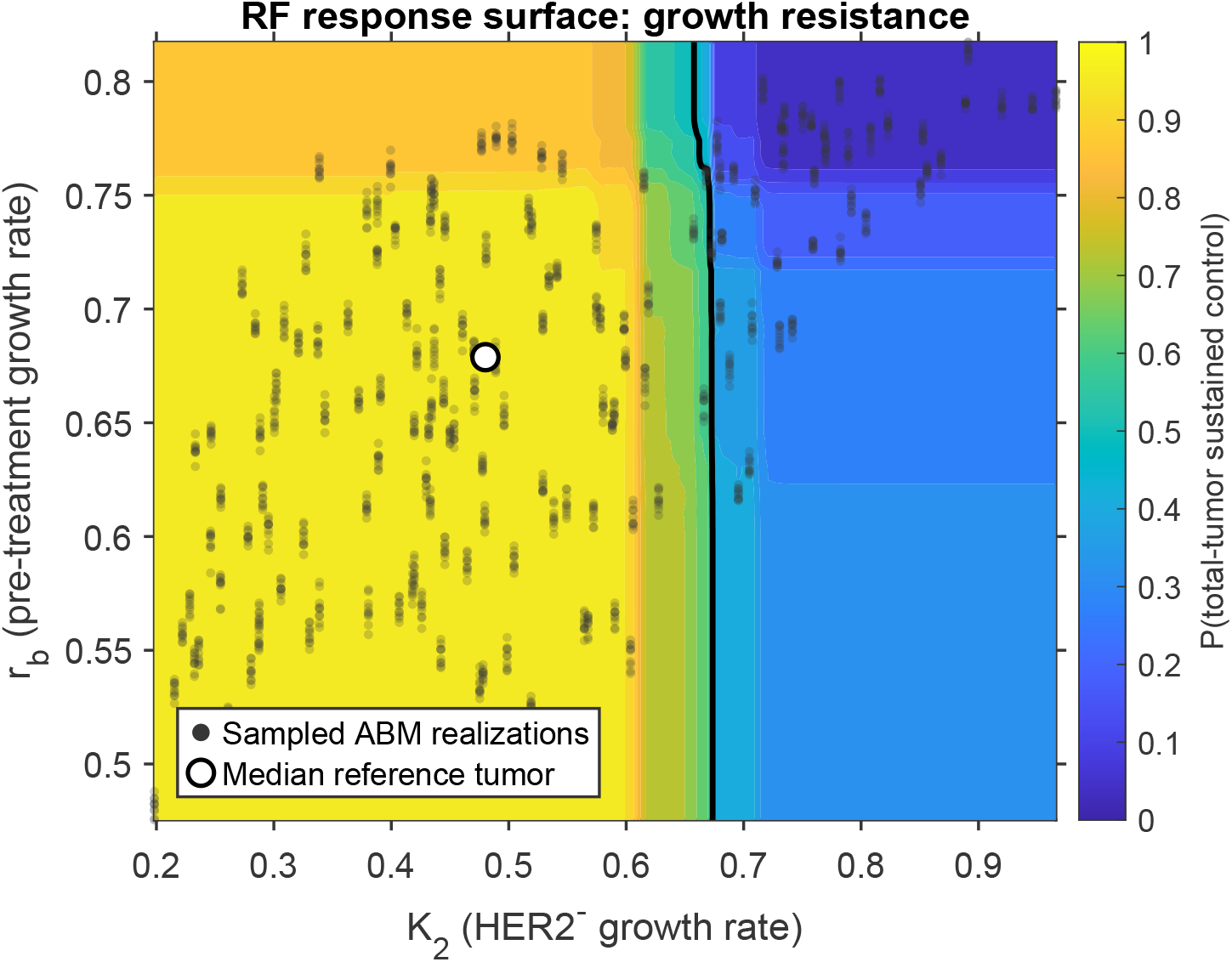
Conditional Random Forest response surface for sustained total-tumor control under standard-dose combination therapy. Color denotes the predicted probability of sustained control as a function of the HER2*^−^* proliferation rate *K*_2_ and the effective pre-treatment growth rate *r_b_*. Sustained control was defined as *N* (10) *≤* 0.25*N* (3), corresponding to a reduction of at least 75% relative to tumor burden at treatment initiation. Predictions were generated using the combined Random Forest model. Non-plotted predictors were fixed at their development-set median values, while *K*_1_ was adjusted with *K*_2_ to preserve the sampled growth-rate relationship. Dark semi-transparent points denote the actual *K*_2_–*r_b_* combinations represented in the development dataset used to fit the Random Forest, illustrating the region of simulation support for the response surface. The white circle denotes the median reference tumor, and the black contour marks a predicted control probability of 0.5.

### 4.9 Illustrative held-out simulated tumor profiles: ABM vs RF

To provide an illustrative comparison between direct agent-based model (ABM) outcomes and Random Forest (RF) predictions, we examined six simulated tumor parameter sets from the final held-out test partition. Each profile corresponds to a distinct tumor parameter set characterized by intrinsic biological properties and early pre-treatment dynamics. Here, “profile” refers only to a simulated tumor parameter set and does not represent a real clinical patient. Three selected parameter sets exhibited stable ABM failure and three exhibited stable ABM success, where all ten stochastic ABM realizations for a given parameter set yielded the same binary treatment outcome. These six cases were selected to illustrate clearly separated low-and high-response regimes and were not used to estimate RF generalization performance.

For each profile, treatment outcome under the fixed combination-therapy protocol was evaluated independently using direct ABM simulations and combined-model RF predictions based on intrinsic parameters and pre-treatment features. Tumor control was assessed at the evaluation horizon *T*_eval_ using the same binary outcome criterion in both frameworks. Specifically, sustained control was defined using the primary criterion *N* (10) *≤* 0.25*N* (3). To account for stochasticity in the ABM, ten stochastic realizations were evaluated for each profile, yielding an empirical ABM success rate. In contrast, the RF surrogate produces a continuous success score for each corresponding held-out realization.

Figure 12 summarizes the resulting illustrative held-out comparison. The left panel shows the mean combined-model RF success score for each profile plotted against the corresponding ABM success rate across its ten stochastic realizations. Because only stable ABM regimes were selected for this illustrative comparison, the ABM success rate is either zero or one. For these selected cases, stable ABM failures are associated with low RF scores, whereas stable ABM successes are associated with higher RF scores. Importantly, this comparison is not intended to demonstrate probabilistic calibration, or to provide an independent estimate of RF predictive performance.

**Figure 12:**
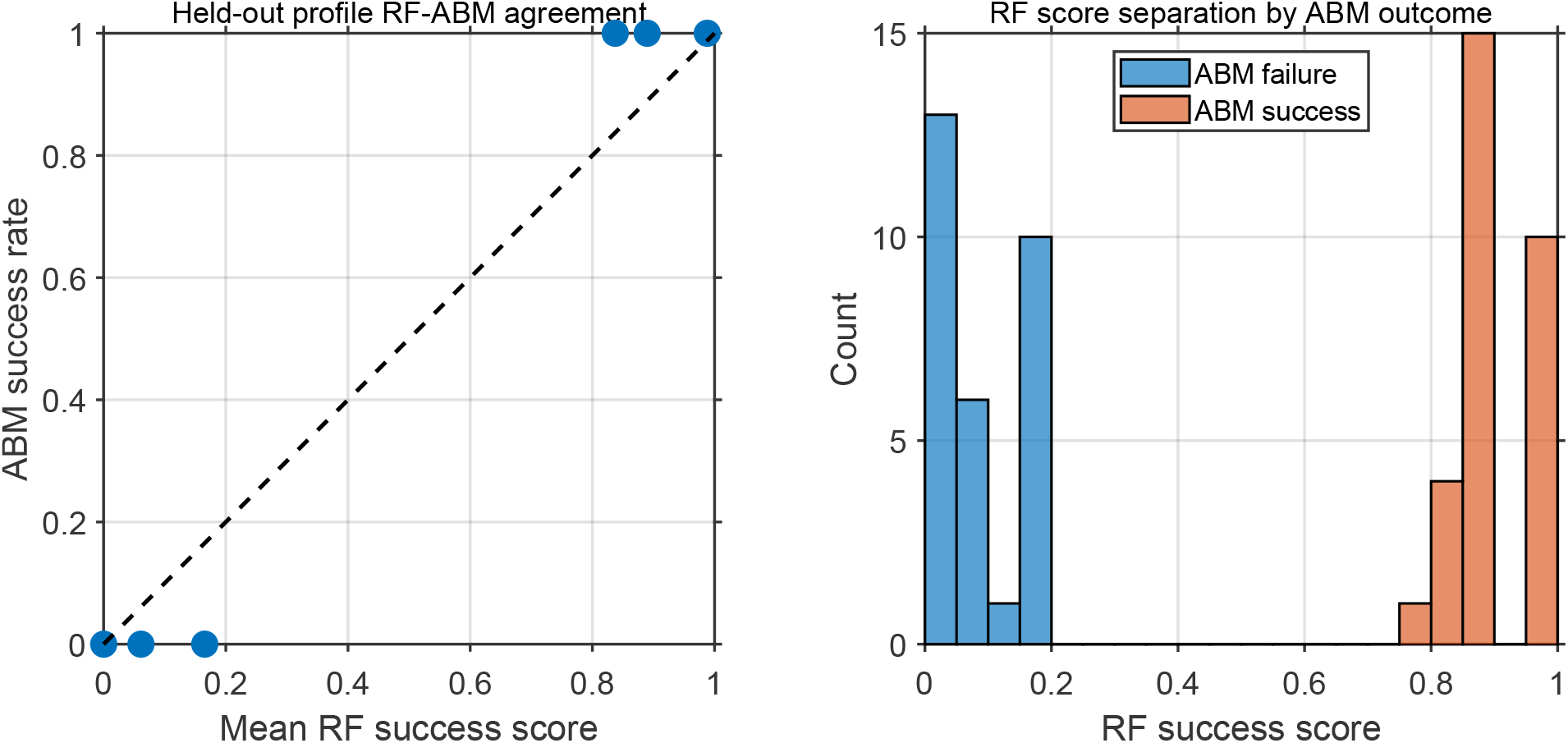
Agreement between Random Forest (RF) predictions and agent-based model (ABM) outcomes for illustrative held-out simulated tumor profiles. Here, each profile represents a simulated tumor parameter set from the final held-out test partition and does not correspond to a real clinical patient. **Left:** Mean RF success score versus ABM success rate for six illustrative held-out profiles, comprising three stable ABM failures and three stable ABM successes. Each profile was evaluated using ten stochastic ABM realizations, with all ten realizations yielding the same binary outcome within that parameter set. The dashed line indicates the identity line (*y* = *x*) as a visual reference. **Right:** Distribution of RF success scores across the 60 stochastic realizations from the same six held-out profiles, separated by ABM failure and ABM success. These six profiles are presented as illustrative examples and are not used to provide an independent estimate of RF predictive performance; quantitative RF generalization was evaluated using all 40 held-out parameter sets, comprising 400 stochastic realizations, as reported in Fig. 9.

**Figure 13:**
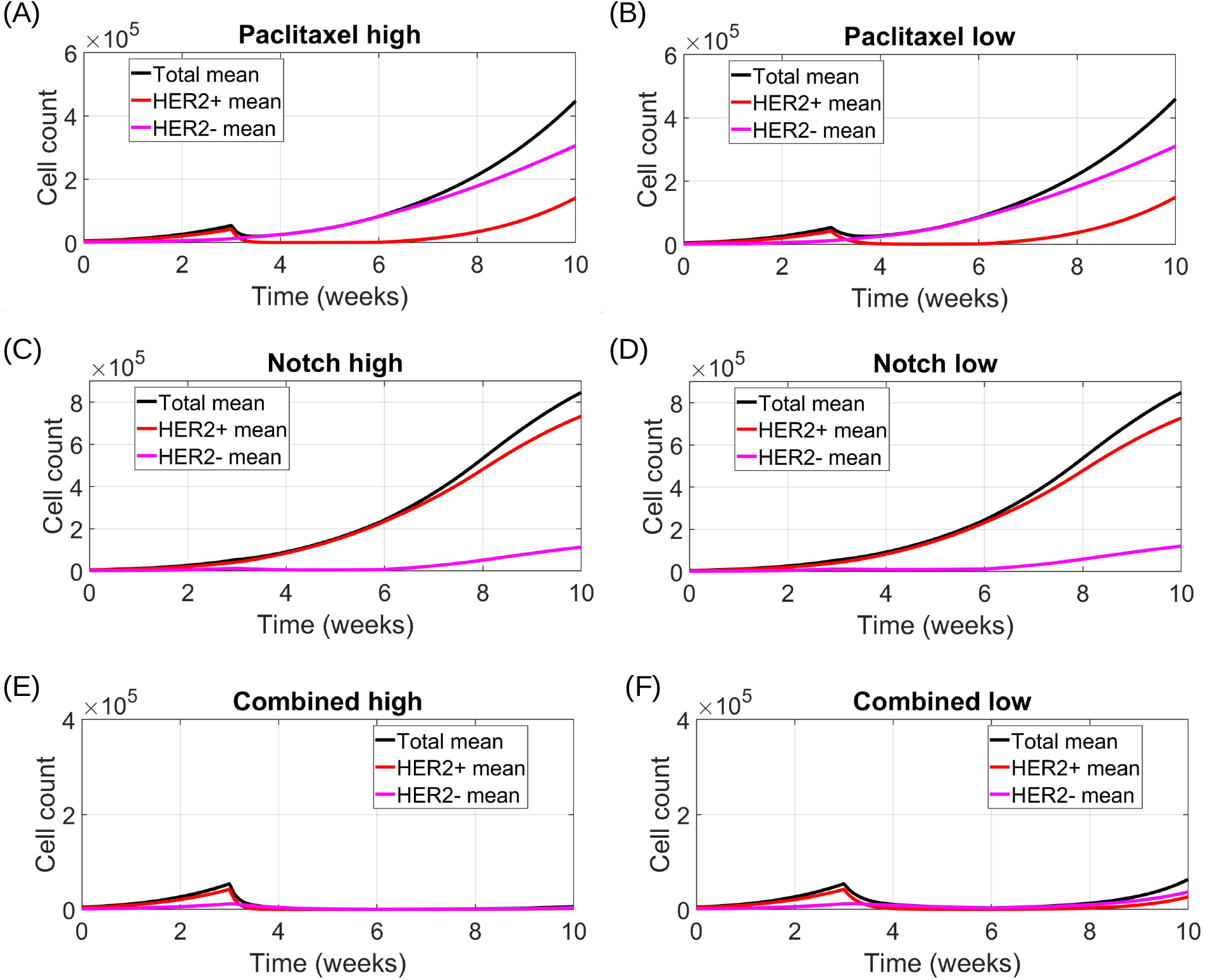
Dose dependence of monotherapy and combination treatment outcomes in the agent-based model. Mean population trajectories over *n* = 100 stochastic realizations are shown for total tumor burden (black), HER2^+^ cells (red), and HER2*^−^* cells (magenta) under high- and low-dose treatment conditions. Shaded regions indicate 95% confidence intervals across stochastic realizations; these intervals are narrow relative to the plotting scale. (A–B) **Paclitaxel monotherapy** selectively suppresses the HER2^+^ population during the treatment window (weeks 3–6), but tumor regrowth occurs after treatment cessation, driven primarily by HER2*^−^* cells. (C–D) **Notch inhibitor monotherapy** preferentially suppresses HER2*^−^* cells, allowing HER2^+^ cells to dominate tumor growth at both dose levels. (E–F) **Combination therapy** exhibits strong dose dependence: high-dose treatment leads to rapid tumor collapse and sustained suppression through the evaluation horizon, whereas low-dose treatment substantially reduces tumor burden during therapy but permits partial regrowth at later times.

**Figure 14:**
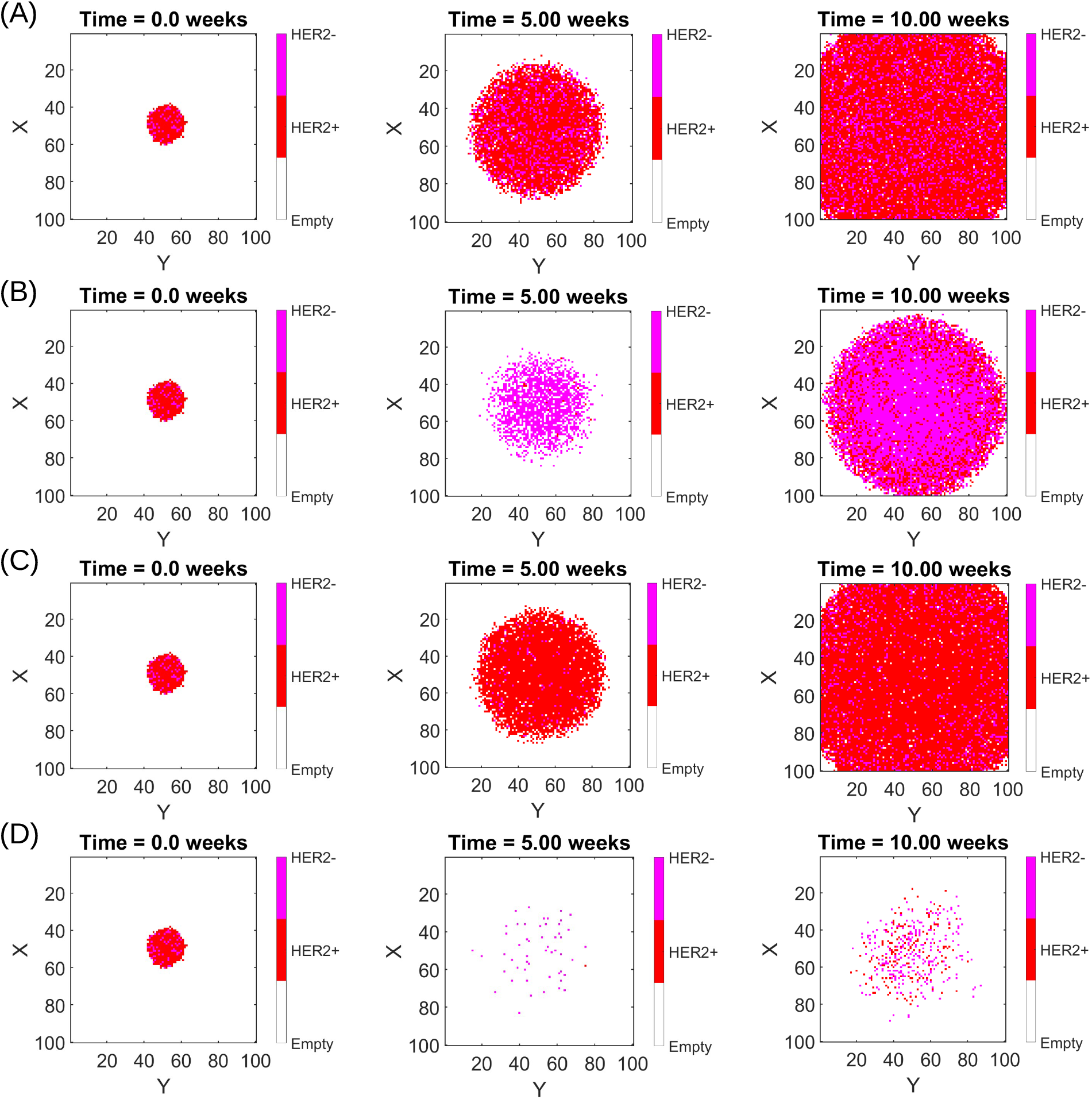
Representative two-dimensional spatial slices of tumor evolution under different treatment strategies. Rows correspond to treatment scenarios: (A) no treatment, (B) Paclitaxel monotherapy, (C) Notch inhibitor monotherapy, and (D) combination therapy. Columns show snapshots at three time points: left, *t* = 0 weeks (initial condition); middle, *t* = 5 weeks; and right, *t* = 10 weeks. Red points denote HER2^+^ tumor cells, magenta points denote HER2*^−^* tumor cells, and white regions correspond to empty lattice sites.

**Figure 15:**
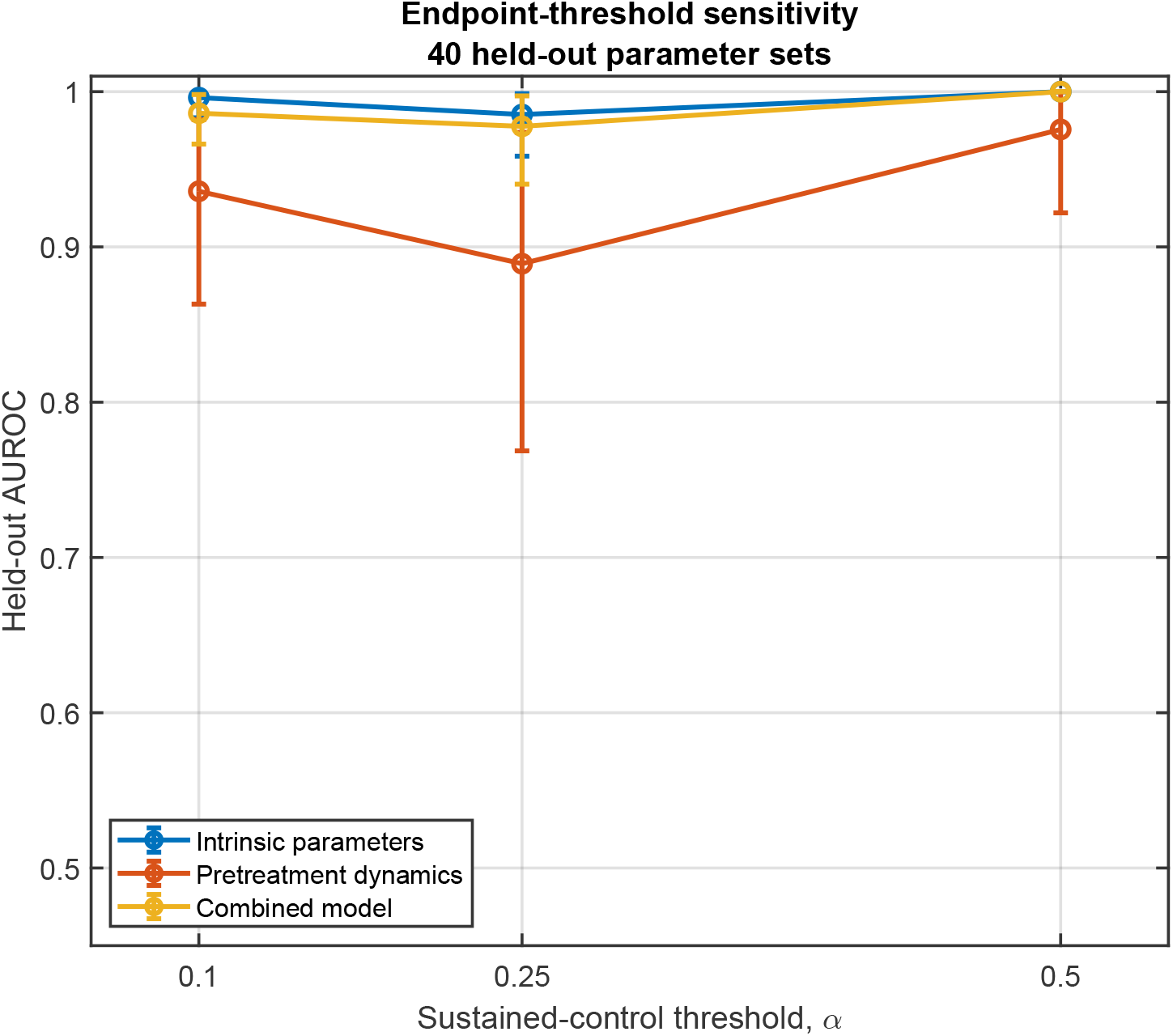
Sensitivity of held-out Random Forest performance to the sustained-control threshold. Held-out AUROC is shown for the intrinsic-parameter, pre-treatment-dynamics, and combined models using *α* = 0.10, 0.25, and 0.50, where sustained total-tumor control was defined as *N* (10) *≤ αN* (3). Each model was evaluated using the same 40 held-out parameter sets, comprising 400 stochastic ABM realizations. Error bars indicate 95% confidence intervals obtained by bootstrap resampling complete held-out parameter-set blocks. The primary analysis used *α* = 0.25.

The right panel further illustrates the comparison at the run level by showing the distribution of RF success scores stratified by ABM outcome. RF scores associated with ABM failures cluster near zero, whereas scores corresponding to ABM successes cluster at substantially higher values, with minimal overlap between the two distributions. Because these six cases were selected for illustration, this separation is interpreted descriptively rather than as an independent validation of the RF.

Together, these results provide an illustrative comparison of RF scores and direct ABM outcomes in selected stable held-out regimes. Quantitative RF generalization was evaluated separately using all 40 held-out parameter sets, comprising 400 stochastic realizations, as reported in Fig. 9.

## 5 Discussion

In this work, we developed an integrated mechanistic and data-driven modeling framework to investigate the role of HER2-mediated phenotypic plasticity in breast cancer treatment response. By combining a spatially resolved agent-based model (ABM) with an interpretable Random Forest (RF) surrogate, we linked single-cell lineage dynamics, spatial organization, and population-level outcomes to predictors of long-term tumor control. This framework enables systematic analysis of how intrinsic tumor heterogeneity, competitive growth asymmetries, and early stochastic dynamics influence therapeutic outcomes across heterogeneous tumor parameter regimes.

The controlled spatial-configuration experiment identifies a treatment-dependent effect of HER2*^−^* localization that cannot be resolved by a well-mixed population model. When tumor geometry, total burden, and phenotype counts were held fixed at treatment initiation, peripheral enrichment of the comparatively Paclitaxel-insensitive HER2*^−^* population substantially increased final tumor burden under Paclitaxel monotherapy. In contrast, peripheral enrichment did not confer a general growth advantage in the untreated condition, and the spatial effect was almost completely eliminated by combination therapy. These results indicate that the therapeutic consequence of phenotype localization depends on the selective pressure imposed by treatment rather than on spatial position alone.

Because the matched tumors correspond to identical HER2^+^ and HER2*^−^* population counts in the ODE formulation, the distinct Paclitaxel responses constitute a specifically spatial prediction of the ABM. The exposed-cell enrichment analysis suggests that comparatively Paclitaxel-insensitive HER2^−^ cells positioned near exposed sites are better able to exploit locally available space following selective depletion of HER2^+^ cells. Simultaneous targeting of both phenotypes removes this opportunity and therefore suppresses the spatial advantage of peripheral HER2*^−^* enrichment. Although this prediction requires experimental validation, it provides a testable hypothesis linking pretreatment phenotype organization to monotherapy escape.

Our results highlight the limitations of monotherapies that selectively target a single phenotype in HER2-heterogeneous tumors. Both paclitaxel and Notch inhibition produce compensatory ecological responses in which suppression of one phenotypic compartment permits expansion of the other. These dynamics are consistent with experimentally observed reversible HER2 state transitions and illustrate how phenotypic plasticity can undermine single-agent therapy. In contrast, combination therapy targeting both HER2-positive and HER2-negative populations reduces this compensatory expansion and can achieve sustained tumor suppression across a wide range of simulated tumor parameter sets. At the same time, the simulations also demonstrate that combination therapy is not universally successful, with treatment failure arising in regimes where intrinsic growth advantages dominate treatment effects.

Importantly, the success of combination therapy is not guaranteed by the model structure alone. Across the simulated parameter ensemble, treatment failure still occurs in regimes where intrinsic proliferation advantages outweigh therapy-induced suppression. In these regimes, even simultaneous targeting of both phenotypes cannot fully counteract rapid population expansion. This highlights that therapeutic outcome is governed by a balance between baseline ecological fitness and treatment pressure, rather than by therapy structure alone.

Because the reference treatment parameters were selected through phenomenological calibration that incorporated the qualitative expectation of stronger suppression under combined treatment, the baseline superiority of combination therapy is not interpreted as an independent prediction of the model. Instead, our analysis focuses on the parameter regimes in which combination therapy succeeds or fails, the determinants of this variation, and the specifically spatial effects that emerge when phenotype abundance is held fixed.

The additional treatment-mechanism sensitivity analysis further clarifies the role of assumptions governing therapy-induced phenotypic plasticity. Variation in the Paclitaxel-associated HER2^+^ *→*HER2*^−^* switching rate produced appreciable changes in residual tumor burden, whereas comparable perturbations of the HER2*^−^ →*HER2^+^ switching rate during Notch inhibition had substantially smaller effects (Appendix Fig. 16). Thus, the magnitude of Paclitaxel-associated phenotypic redirection represents an influential model assumption and should not be interpreted as independently identified from the available data. Importantly, substantial tumor suppression was retained across all switching scenarios examined.

**Figure 16:**
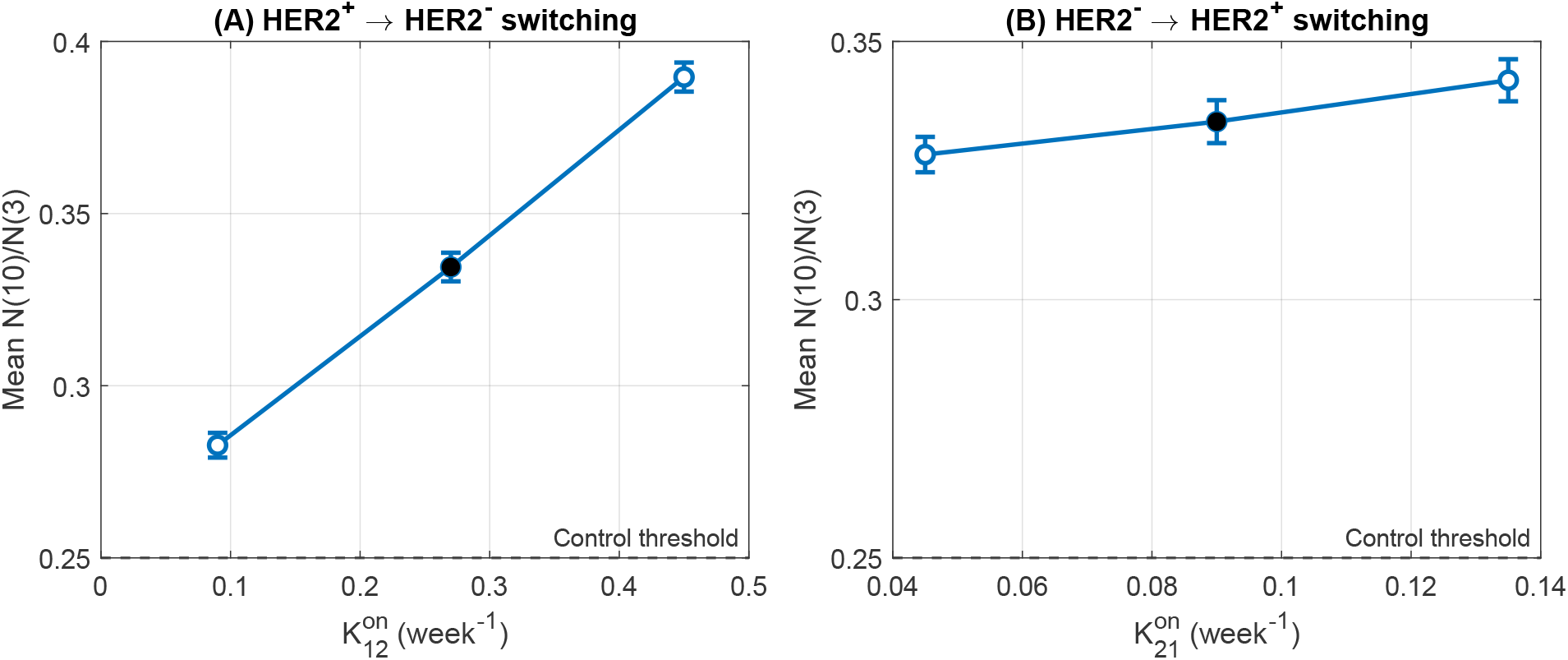
Sensitivity of combination-therapy response to treatment-modulated phenotypic switching assumptions. Mean normalized week-10 tumor burden, *N* (10)*/N* (3), is shown under standard-dose combination therapy for variation in (A) the HER2^+^ *→*HER2*^−^* asymmetric switching rate during Paclitaxel exposure, *K*^on^, and (B) the HER2*^−^ →*HER2^+^ asymmetric switching rate during Notch inhibition, *K*^on^. Each condition contains *n* = 100 paired stochastic simulations. Error bars indicate 95% confidence intervals for the mean. Filled black circles denote the reference treatment assumptions, *K*^on^ = 0.27 week*^−^*^1^ and *K*^on^ = 0.09 week*^−^*^1^. The horizontal dashed line denotes the primary sustained-control threshold, *N* (10)*/N* (3) = 0.25. All other treatment-modified parameters were held at their standard values.

A central contribution of this study is clarification of the modeling hierarchy between deterministic and spatially resolved approaches. Population-level ODE models accurately predict long-term phenotypic equilibria in the large-population limit and provide an important theoretical baseline. However, such models necessarily average over spatial structure and stochastic lineage effects. Our results suggest that early founder variability, variance in small colonies, and localized spatial rearrangements can alter transient treatment trajectories in ways that are not captured by well-mixed formulations. The ABM provides a framework for resolving these mechanisms explicitly and for connecting population-level dynamics to spatially structured cell interactions.

The RF surrogate complements the mechanistic model by providing a compact and interpretable summary of high-dimensional simulation outcomes. Trained on ensemble ABM simulations and restricted to pre-treatment and early-trajectory features, the RF accurately predicts long-term treatment outcomes for parameter combinations not used during training. Because the training and testing sets were separated at the parameter level, this evaluation measures interpolation across previously unseen biological parameter combinations within the sampled domain rather than extrapolation beyond it. The high held-out discrimination should be interpreted in the context of this simulation-surrogate task: the RF predicts ABM-generated outcomes from intrinsic parameters and pre-treatment features that directly encode the tumor dynamics governing those simulated responses. Thus, the high AU-ROC reflects a structured response boundary within the sampled model domain rather than clinical predictive accuracy.

The grouped permutation-importance and conditional response-surface analyses reveal a clear growth-dominated transition in treatment response under combination therapy. In particular, the intrinsic proliferation rate of the HER2-negative population and the effective pre-treatment growth dynamics define a boundary separating regimes of sustained suppression from regimes of tumor escape, whereas phenotype switching rates exert a comparatively weaker influence on treatment outcome. These findings suggest that baseline competitive fitness and net expansion dynamics can play a larger role in determining therapeutic success than fine-scale modulation of switching behavior.

More broadly, the experimental lineage and colony comparisons used in this study provide biological benchmarking of selected model behaviors but do not constitute independent validation of the full ABM. Validation against independent experimental datasets, particularly datasets that jointly resolve HER2 phenotype, spatial organization, and treatment response, remains an important direction for future work.

### Modeling assumptions on phenotypic switching

In the present model, transitions between HER2^+^ and HER2*^−^* states are implemented through asymmetric division events. This formulation reflects the lineage-based phenotypic plasticity observed in experimental studies of circulating breast cancer cells, where colonies derived from single cells generate mixed HER2^+^ and HER2*^−^* populations during expansion [7]. While these experiments demonstrate reversible phenotypic transitions, the precise intracellular timing of switching events remains unresolved. Coupling switching to division therefore provides a minimal representation of reversible phenotypic inheritance while avoiding additional poorly constrained parameters. Future work could explore alternative formulations allowing continuous stochastic switching or therapy-induced transitions independent of division.

Beyond the specific therapies considered here, the modeling framework is readily extensible. Incorporating additional components of the tumor microenvironment, including cytotoxic T cells, immune suppression, and cytokine signaling, would allow investigation of how immune-mediated selection pressures interact with HER2 plasticity and spatial structure. Extending the ABM to model cell-based immunotherapies such as CAR-T treatment could further enable exploration of antigen heterogeneity, trafficking constraints, and tumor–immune co-evolution in solid tumors.

Future work may also explore alternative treatment schedules and adaptive strategies. While the present study focused on fixed monotherapy and combination protocols, the ABM–RF framework provides a foundation for evaluating sequential, dose-modulated, or control-optimized regimens designed to exploit tumor ecological vulnerabilities. Coupling the surrogate model with optimization or reinforcement learning approaches may enable systematic identification of treatment strategies that maintain tumors in controllable states while delaying resistance.

In summary, this study demonstrates that stochastic lineage dynamics, phenotype localization, heterogeneous tumor growth, and competitive growth asymmetries can influence therapy response in HER2-heterogeneous breast cancer. In particular, phenotype localization can alter treatment response even when tumor burden and phenotype abundance are held fixed, revealing a specifically spatial effect that is absent from the corresponding well-mixed population model. By integrating spatially resolved mechanistic modeling with interpretable machine learning, this framework provides a scalable approach for analyzing simulated tumor dynamics and identifying growth-driven resistance regimes that shape therapeutic outcomes.

## 6 Data Availability

The MATLAB code, simulated data, analysis outputs, and figure-generation scripts used to reproduce the analyses reported in this study are publicly available on GitHub.

## **A** Model Details and Parameters

This appendix provides full details of the agent-based model update rules and the parameter definitions used in all simulations. The present model considers tumor-cell–intrinsic dynamics only and does not include immune or stromal components.

### **A.1** Tumor Cell Update Rules

At each discrete time step of length *dt*, tumor cells undergo stochastic events modeled as Poisson processes with phenotype-specific rates.

- **Division:** Each HER2^+^ (type A) and HER2^−^ (type B) tumor cell attempts to divide with probability determined by its division rate. Division may be symmetric, producing two daughter cells of the same phenotype, or asymmetric, resulting in phenotypic switching. Asymmetric division from HER2^+^ to HER2^−^ occurs at rate *K*_12_, while conversion from HER2^−^ to HER2^+^ occurs at rate *K*_21_.
- **Movement:** Each tumor cell attempts to move to a randomly chosen neighboring lattice site with probability *µ dt*. Movement occurs only if the target site is unoccupied.
- **Death:** Each tumor cell undergoes spontaneous death with probability *δ dt*, after which the cell is removed from the lattice.

### **A.2** Drug Effects

Therapeutic interventions are implemented as time-dependent modifications of tumor-cell division, phenotypic switching, and death rates during prescribed treatment windows.

- **Paclitaxel:** During Paclitaxel treatment, the division rate of HER2^+^ tumor cells is reduced and their death rate is increased to represent cytotoxic chemotherapy effects. In addition, the rate of asymmetric division from HER2^+^ to HER2^−^ is increased, modeling stress-induced phenotypic transitions. All rates associated with HER2^−^ cells remain unchanged during Paclitaxel exposure.
- **Notch Inhibitor:** During Notch inhibitor treatment, the division rate of HER2^−^ tumor cells is reduced and their death rate is increased, reflecting selective loss of Notch-dependent survival and proliferative capacity. The rate of phenotypic conversion from HER2^−^ to HER2^+^ is unchanged, preserving phenotypic plasticity during treatment.

### **A.3** Implementation of matched spatial configurations

The controlled spatial configurations were constructed from a single reference tumor generated by untreated simulation from week 0 to week 3. In the realization used for this experiment, the common week-3 tumor contained 54,535 cells, comprising 42,458 HER2^+^ cells and 12,077 HER2*^−^* cells. The complete week-3 lattice was copied before phenotype redistribution so that the occupied-site mask, total tumor burden, and phenotype counts were identical across all spatial configurations, which were derived from the same pretreatment tumor realization.

Let *O* denote the set of occupied lattice sites in the reference tumor. The tumor centroid was calculated as

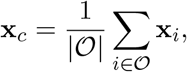

where **x***_i_* is the three-dimensional coordinate of occupied site *i*. The occupied sites were then ranked according to their Euclidean distance

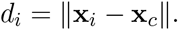

For the centrally enriched configuration, the prescribed number of HER2*^−^* labels was assigned to the occupied sites with the smallest values of *d_i_*. For the peripherally enriched configuration, HER2*^−^* labels were assigned to the occupied sites with the largest values of *d_i_*. For the randomly distributed configuration, the same number of HER2*^−^* labels was assigned to a random permutation of the occupied sites. In each case, all remaining occupied sites were assigned the HER2^+^ phenotype. No occupied sites were added, removed, or moved during construction of the matched configurations.

Exposed tumor cells were identified using the three-dimensional 26-site Moore neighborhood. Computationally, the occupied tumor mask was convolved with a 3 *×* 3 *×* 3 kernel containing ones at all neighboring positions and zero at its central position. An occupied site with fewer than 26 occupied neighbors was classified as exposed. This definition includes cells adjacent to the external tumor surface as well as cells adjacent to internal unoccupied lattice sites. The exposed-cell enrichment index was then calculated as the HER2*^−^* fraction among exposed cells divided by the global HER2*^−^* fraction.

Following construction, each matched grid was used as the initial condition at week 3 for untreated, Paclitaxel, and combination-therapy continuations. Within each treatment condition, corresponding replicate numbers used the same continuation seed for the central, random, and peripheral configurations. Phenotype labels were not fixed after treatment initiation; cells subsequently divided, migrated, switched phenotype, and died according to the standard agent-based model rules.

### **A.4** Baseline Model Parameters

### A.5 Treatment-Dependent Parameter Modifications

The treatment-dependent event rates were selected jointly through manual phenomenological calibration rather than estimated independently. The reference parameter set was chosen to reproduce the experimentally motivated qualitative response hierarchy of continued tumor growth in the untreated condition, incomplete suppression under either monotherapy, and the strongest tumor suppression under simultaneous paclitaxel and Notch inhibition. These values defined the standard treatment condition and were held fixed unless treatment intensity was explicitly varied.

Because this phenomenological calibration included the qualitative expectation of stronger suppression under combined treatment, the baseline superiority of combination therapy is not interpreted here as an independently predicted result. Instead, the analyses focus on the parameter regimes in which combination therapy succeeds or fails, the determinants of that variation, and the specifically spatial effects that emerge when phenotype abundance is held fixed.

When treatment intensity was varied, each treatment-modified event rate was scaled relative to its baseline value according to

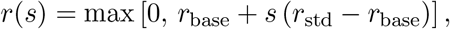

where *r*_base_ is the untreated baseline rate, *r*_std_ is the standard-treatment rate reported in Table 2, and *s* is the dose-scaling factor. Thus, *s* = 1 corresponds to the standard treatment condition, while *s* = 0.75 and *s* = 1.25 represent the low- and high-dose conditions, respectively. The nonnegativity constraint prevents dose scaling from producing invalid negative stochastic event rates. Parameters not modified by treatment remained at their baseline values.

### **A.6** Treatment-mechanism sensitivity analysis

To evaluate the robustness of the therapy-dependent phenotypic-switching assumptions, we performed an additional sensitivity analysis under standard-dose combination therapy (*s* = 1). All direct treatment effects on phenotype-specific proliferation and death rates were retained at their reference values in Table 2, while the treatment-period asymmetric switching rates were varied independently. For the Paclitaxel-associated HER2^+^ *→*HER2*^−^* transition, we considered *K*^on^ *∈ {*0.09, 0.27, 0.45*}* week*^−^*^1^, where 0.09 corresponds to no treatment-induced increase above the untreated switching rate, 0.27 is the reference treatment value, and 0.45 represents a stronger treatment-induced transition. For the HER2*^−^ →*HER2^+^ transition during Notch inhibition, we considered *K*^on^ *∈ {*0.045, 0.09, 0.135*}* week*^−^*^1^, with 0.09 corresponding to the reference assumption that this transition remains unchanged during treatment.

Because the reference condition is shared between the two sensitivity series, five unique treatment conditions were simulated. Each condition was evaluated using *n* = 100 stochastic realizations, giving 500 simulations in total. A paired-seed design was used: for each replicate, the same random seed was reset before each treatment condition so that all five conditions shared an identical pretreatment realization through week 3. Combination therapy was applied continuously from week 3 to week 6, and tumors were followed through week 10. Sensitivity was summarized using the normalized final tumor burden,

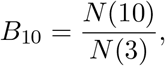

and paired differences in *B*_10_ relative to the reference treatment condition. Mean values and 95% confidence intervals were calculated across the 100 stochastic replicates.

## **B** Additional Results: Dose Dependence and Spatial Organization

To assess robustness with respect to treatment strength, we examined low- and high-dose variants of mono- and combination therapies using the agent-based model.

Fig 13 shows that increasing dose primarily enhances tumor control under combination therapy, while monotherapies remain insufficient to prevent long-term tumor persistence across dose levels.

To complement the population-level dynamics presented in the main text, we next examine the spatial organization of tumor cells under different treatment strategies.

Fig 14 illustrates how treatment reshapes tumor spatial organization beyond changes in total burden. Without treatment, the tumor grows outward from the initial cluster in a largely continuous manner, with HER2^+^ and HER2*^−^* cells remaining intermingled. Under Paclitaxel monotherapy, HER2^+^ cells are progressively depleted and HER2*^−^* cells come to dominate the spatial structure, filling regions left vacant by therapy-induced cell death. In contrast, Notch inhibition selectively suppresses HER2*^−^* cells, allowing HER2^+^ cells to spread and form a more uniform spatial population.

The most striking behavior occurs under combination therapy. Rather than shrinking uniformly, the tumor fragments into smaller, scattered clusters, and neither phenotype establishes clear spatial dominance. Even at later times, these fragmented clusters persist, indicating that simultaneously targeting both phenotypes disrupts the tumor’s ability to maintain a coherent spatial structure. Overall, these spatial visualizations highlight treatment-induced changes in tumor organization that are not apparent from population-level curves alone.

### **B.1** Random Forest endpoint-threshold sensitivity

To assess whether Random Forest performance depended strongly on the specific definition of sustained tumor control, we repeated the classification analysis using three endpoint thresholds, *α ∈ {*0.10, 0.25, 0.50*}*, where sustained control was defined as *N* (10) *≤ αN* (3). The same development and held-out parameter-set partitions were retained for each analysis. The intrinsic-parameter and combined models maintained very high held-out discrimination across all three thresholds, whereas the pre-treatment-dynamics model remained predictive but showed greater sensitivity to the endpoint definition. These results indicate that the principal RF conclusions were not driven by the primary choice *α* = 0.25.

### **B.2** Sensitivity to treatment-modulated phenotypic switching

The treatment response was substantially more sensitive to the assumed Paclitaxel-associated HER2^+^ *→*HER2*^−^* switching rate than to the reverse switching rate during Notch inhibition (Fig. 16). At the reference value *K*^on^ = 0.27 week*^−^*^1^, the mean normalized week-10 tumor burden was *N* (10)*/N* (3) = 0.334. Eliminating the treatment-induced increase in HER2^+^ *→*HER2*^−^* switching (*K*^on^ = 0.09 week*^−^*^1^) reduced the mean normalized burden to 0.283, with a paired difference from the reference condition of *−*0.052 (95% CI: *−*0.057 to *−*0.047). In contrast, increasing *K*^on^ to 0.45 week*^−^*^1^ increased the mean normalized burden to 0.390, corresponding to a paired difference of 0.055 (95% CI: 0.050 to 0.061).

Perturbations of the HER2*^−^ →*HER2^+^ switching rate during Notch inhibition produced substantially smaller changes. Reducing *K*^on^ from the reference value of 0.09 to 0.045 week*^−^*^1^ changed the mean normalized week-10 burden from 0.334 to 0.328, with a paired difference of *−*0.006 (95% CI: *−*0.011 to *−*0.002). Increasing *K*^on^ to 0.135 week*^−^*^1^ increased the mean burden to 0.342, with a paired difference of 0.008 (95% CI: 0.004 to 0.012). Thus, residual tumor burden was quantitatively sensitive to the magnitude of Paclitaxel-associated HER2^+^ *→*HER2*^−^* switching, whereas moderate variation in the reverse transition during Notch inhibition had a comparatively small effect. Nevertheless, mean week-10 tumor burden remained below 40% of the week-3 burden across all tested switching assumptions.

## References

[1] D. J. Slamon, G. M. Clark, S. G. Wong, W. J. Levin, A. Ullrich, and W. L. McGuire, “Studies of the HER-2/neu proto-oncogene in human breast and ovarian cancer,” Science, vol. 235, no. 4785, pp. 177–182, 1987.

[2] S. M. Swain, J. Baselga, S.-B. Kim, J. Ro, V. Semiglazov, M. Campone, E. Ciruelos, J.-M. Ferrero, A. Schneeweiss, S. Heeson, E. Clark, G. Ross, M. C. Benyunes, and J. Cortés, “Pertuzumab, trastuzumab, and docetaxel in her2-positive metastatic breast cancer,” The New England journal of medicine, vol. 372, no. 8, pp. 724–734, 2015.

[3] D. J. Slamon et al., “Use of chemotherapy plus a monoclonal antibody against her2 for metastatic breast cancer that overexpresses her2,” New England Journal of Medicine, vol. 344, no. 11, pp. 783–792, 2001.

[4] J. Baselga et al., “Pertuzumab plus trastuzumab plus docetaxel for metastatic breast cancer,” New England Journal of Medicine, vol. 366, no. 2, pp. 109–119, 2012.

[5] S. Modi et al., “Trastuzumab deruxtecan in previously treated her2-low advanced breast cancer,” New England Journal of Medicine, vol. 387, no. 1, pp. 9–20, 2022.

[6] P. Tarantino et al., “Her2-low breast cancer: Pathological and clinical landscape,” Journal of Clinical Oncology, vol. 38, no. 17, pp. 1951–1962, 2020.

[7] N. V. Jordan, A. Bardia, B. S. Wittner, C. Benes, M. Ligorio, Y. Zheng, M. Yu, T. K. Sundaresan, J. A. Licausi, R. Desai, R. M. O’Keefe, R. Y. Ebright, M. Boukhali, S. Sil, M. L. Onozato, A. J. Iafrate, R. Kapur, D. Sgroi, D. T. Ting, M. Toner, S. Ramaswamy, W. Haas, S. Maheswaran, and D. A. Haber, “Her2 expression identifies dynamic functional states within circulating breast cancer cells,” Nature (London), vol. 537, no. 7618, pp. 102–106, 2016.

[8] P. B. Gupta, C. M. Fillmore, G. Jiang, S. D. Shapira, K. Tao, C. Kuperwasser, and …, “Stochastic state transitions give rise to phenotypic equilibrium in populations of cancer cells,” Cell, vol. 146, no. 4, pp. 633–644, 2011.

[9] S. V. Sharma et al., “A chromatin-mediated reversible drug-tolerant state in cancer cell subpopulations,” Cell, vol. 141, no. 1, pp. 69–80, 2010.

[10] X. Li and D. Thirumalai, “A mathematical model for phenotypic heterogeneity in breast cancer with implications for therapeutic strategies,” Journal of the Royal Society interface, vol. 19, no. 186, pp. 20 210 803–, 2022.

[11] C. L. Chaffer, I. Brueckmann, C. Scheel, A. J. Kaestli, P. A. Wiggins, L. O. Rodrigues, M. Brooks, F. Reinhardt, Y. Su, K. Polyak, L. M. Arendt, C. Kuperwasser, B. Bierie, and R. A. Weinberg, “Normal and neoplastic nonstem cells can spontaneously convert to a stem-like state,” Proceedings of the National Academy of Sciences - PNAS, vol. 108, no. 19, pp. 7950–7955, 2011.

[12] C. E. Meacham and S. J. Morrison, “Tumour heterogeneity and cancer cell plasticity,” Nature (London), vol. 501, no. 7467, pp. 328–337, 2013.

[13] S. M. Tovey, S. Brown, J. C. Doughty, E. A. Mallon, T. G. Cooke, and J. Edwards, “Poor survival outcomes in HER2-positive breast cancer patients with low-grade, node-negative tumours,” British Journal of Cancer, vol. 100, pp. 680–683, 2009.

[14] F. Y. Wong, C. S. P. Yip, and E. T. Chua, “Implications of her2 amplification in small, node-negative breast cancers: Do asians differ?” World journal of surgery, vol. 36, no. 2, pp. 287–294, 2012.

[15] A. Gavrilova, T. Jackson, and N. Rahman, “Phenotypic plasticity and competition shape therapy sequencing in her2+/her2– breast cancer: A mathematical framework,” Journal of Theoretical Biology, vol. 632, p. 112533, 2026.

[16] A. Anderson and M. Chaplain, “Continuous and discrete mathematical models of tumor-induced angiogenesis,” Bulletin of mathematical biology, vol. 60, no. 5, pp. 857–899, 1998.

[17] T. L. Jackson and H. M. Byrne, “A mathematical model to study the effects of drug resistance and vasculature on the response of solid tumors to chemotherapy,” Mathematical biosciences, vol. 164, no. 1, pp. 17–38, 2000.

[18] A. R. A. Anderson and V. Quaranta, “Integrative mathematical oncology,” Nature reviews. Cancer, vol. 8, no. 3, pp. 227–234, 2008.

[19] J. Metzcar et al., “A review of cell-based computational modeling in cancer biology,” JCO Clinical Cancer Informatics, vol. 3, pp. 1–13, 2019.

[20] K. M. Storey and T. L. Jackson, “An agent-based model of combination oncolytic viral therapy and anti-pd-1 immunotherapy reveals the importance of spatial location when treating glioblastoma,” Cancers, vol. 13, no. 21, pp. 5314–, 2021.

[21] A. Ghaffarizadeh, R. Heiland, S. H. Friedman, S. M. Mumenthaler, and P. Macklin, “Physicell: An open source physics-based cell simulator for 3-d multicellular systems,” PLoS computational biology, vol. 14, no. 2, pp. e1 005 991–, 2018.

[22] H. Korkaya, A. Paulson, E. Charafe-Jauffret, C. Ginestier, M. Brown, J. Dutcher, S. G. Clouthier, and M. S. Wicha, “Regulation of mammary stem/progenitor cells by PTEN/Akt/*β*-catenin signaling,” Proceedings of the National Academy of Sciences, vol. 106, no. 2, pp. 702–707, 2009.

[23] O. Meurette and P. Mehlen, “Notch signaling in the tumor microenvironment,” Cancer Cell, vol. 34, no. 4, pp. 536–548, 2019.

[24] L. Breiman, “Random forests,” Machine Learning, vol. 45, no. 1, pp. 5–32, 2001.

